# Circadian phase reconfigures information transmission in the mouse visual thalamus

**DOI:** 10.64898/2026.07.24.740496

**Authors:** Abigail Pienaar, Jessica Rodgers, Riccardo Storchi, Annette E Allen

## Abstract

Circadian clocks are a ubiquitous feature of the mammalian brain, yet whether they regulate how neural circuits encode and transmit information remains largely unknown. Here, we address this question in the mouse visual thalamus (dLGN), a key relay between retina and cortex that provides a tractable model for studying sensory information transfer. Using wireless electrophysiological recordings from awake, behaving mice, we show that spontaneous dLGN activity is under circadian control, peaking during the late subjective day. Combining visual stimulation, information-theoretic analyses, and optogenetic interrogation of retinogeniculate transmission, we demonstrate that circadian phase selectively reconfigures the visual code. Although fundamental tuning properties remain largely stable, the impact of circadian phase on information transfer depends on visual scene statistics, promoting a more efficient coding strategy at night under high-contrast conditions. Together, these findings identify circadian clocks as dynamic regulators of neuronal communication that actively reshape sensory information processing across the day-night cycle.

## Introduction

Endogenous circadian clocks provide an internal representation of time-of-day, enabling biological systems to anticipate rhythmic changes in the environment. In mammals, the master circadian pacemaker resides in the hypothalamic suprachiasmatic nucleus (SCN) (Hastings et al., 2018). However, rhythmic patterns in gene expression, neuronal connectivity, neurochemistry, and excitability extend beyond the SCN and are evident throughout the brain (Abe et al., 2002; Guilding and Piggins, 2007; Mendoza, 2025; Paul et al., 2020). While the ultimate outputs of these rhythms are changes in physiological state and animal behaviour, their widespread presence and effects on neuronal excitability suggest a more fundamental role in regulating the operation of neural circuits themselves (Yamashita et al., 2025). Consistent with this idea, circadian modulation has been described at multiple levels of neural organisation, including synaptic connectivity (Azeez et al., 2018; Nowacka et al., 2025; Severin et al., 2024) and sensory function (Andrade, 2021; Basinou et al., 2017; Besharse and McMahon, 2016; Bumgarner et al., 2021; Valdez, 2019). However, there have been remarkably few direct investigations into whether circadian clocks alter information coding or transmission within neural circuits (see (Donen et al., 2025; Moya-Díaz et al., 2022)). As such, whether (and how) circadian clocks alter the encoding and transmission of information within neural circuits remains largely unknown.

The visual thalamus (dorsal lateral geniculate nucleus (dLGN)) provides a powerful system in which to address this question. Vision is profoundly shaped by time-of-day, primarily reflecting the dramatic daily cycle in ambient irradiance arising from the Earth’s axial rotation. In the mammalian retina, circadian clocks drive rhythmic changes in cellular physiology, biochemistry, and morphology (Besharse and McMahon, 2016; Kamphuis et al., 2005; Ruan et al., 2006; Storch et al., 2007; Tosini and Menaker, 1996), but probing information transfer here *in vivo* remains technically challenging. In contrast, the dLGN presents a more tractable target: by combining high-density electrophysiological recordings with precisely controlled visual stimuli, we can quantify multiple dimensions of visual coding *in vivo*, including receptive field structure, tuning properties, information transfer and synaptic gain. Furthermore, dLGN activity is also strongly modulated by behavioural state and ambient irradiance (Aydın et al., 2018; Davis et al., 2015; Erisken et al., 2014; Storchi et al., 2017, 2015), allowing circadian influences on information processing to be interpreted alongside other established modulatory factors.

Using multi-channel electrophysiological recordings in freely moving mice, we show that spontaneous activity within the dLGN is robustly regulated by circadian phase, peaking during the late subjective day independently of locomotor behaviour or ambient illumination. Combining visual stimulation, information-theoretic analyses and optogenetic interrogation of retinogeniculate transmission, we further demonstrate that circadian phase selectively reshapes visual coding within this circuit. While core tuning properties remain largely stable across the day-night cycle, receptive field size, spatial coding, and synaptic gain are systematically reconfigured. These changes are consistent with a shift in coding strategy, favouring greater efficiency at night and higher information capacity during the day. Together, these findings identify circadian time as a fundamental determinant of sensory coding and show that the visual thalamus operates in distinct regimes across the day-night cycle.

## Results

### Circadian phase modulates spontaneous neural activity in dLGN

Recent work has revealed clear circadian rhythmicity in the expression of c-fos, a marker of neural excitability, within the dLGN (Yamashita et al., 2025). Electrophysiological studies performed in anaesthetized animals and *ex-vivo* brain slices have, however, provided mixed evidence as to whether these rhythms translate into time-of-day-dependent changes in neuronal activity (Brown et al., 2011; Chrobok et al., 2021). To address this question, we recorded spontaneous activity from dLGN neurons in awake, freely behaving mice using chronically implanted multichannel electrodes coupled to a wireless headstage (Fig 1A,B). We collected 10-minute recordings of spontaneous multiunit activity together with locomotor behaviour every 2 h across a complete 24-h cycle in animals maintained in constant darkness (Fig 1C).

**Figure 1.**
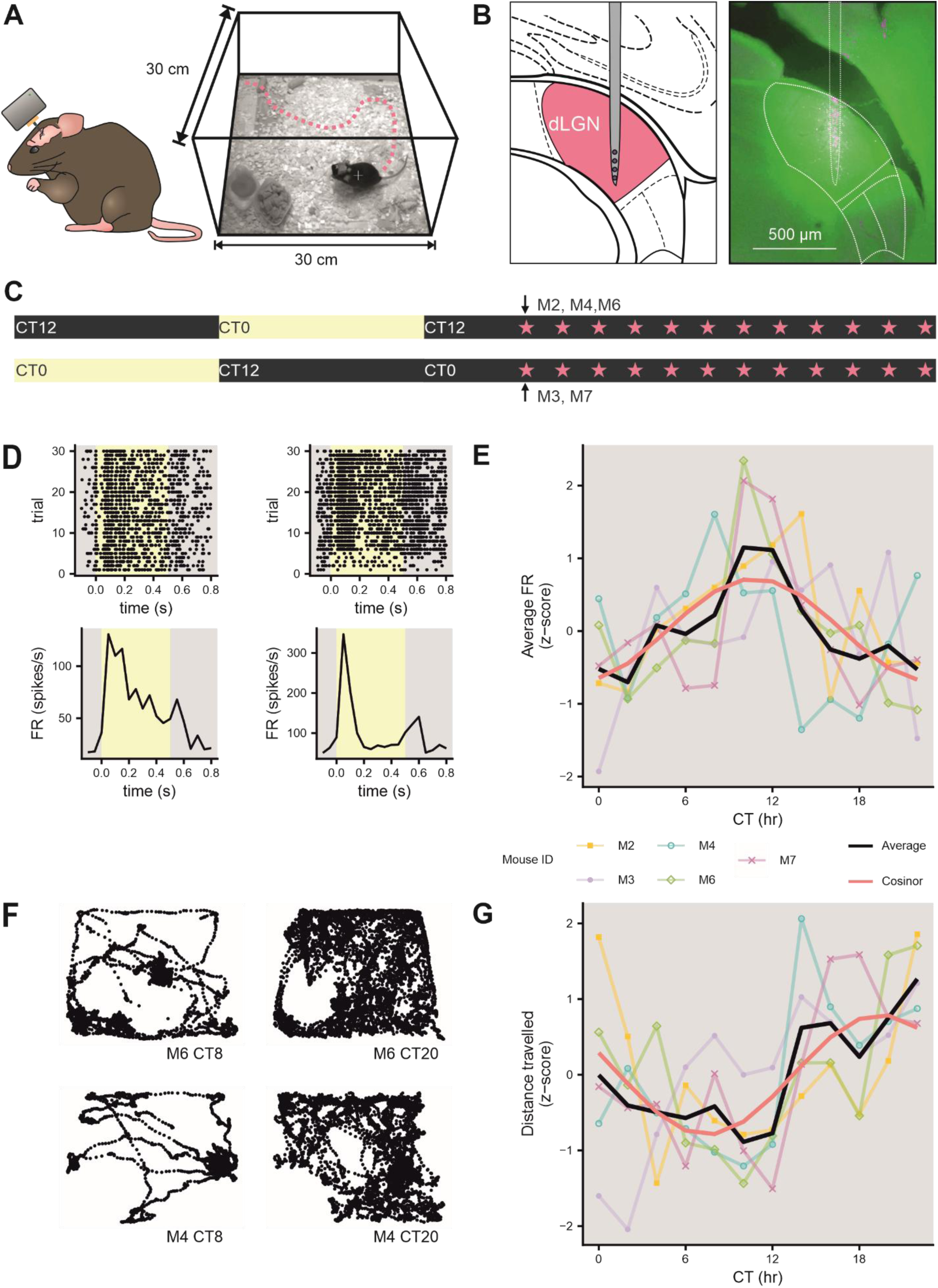
Circadian modulation of spontaneous firing activity in the dLGN of awake mice. 1A. Schematic of wireless headstage used for multiunit electrophysiological recordings in freely behaving mice (left), and image of mouse behavioural arena (right). Mice were maintained in constant darkness and allowed to explore an open-field arena. B. Schematic showing electrode placement within the dLGN and the layout of recording contacts on a single shank (left), together with a representative histological section showing the electrode tract labelled with DiI (right). Only one electrode shank is shown due to sagittal orientation of electrode. C. Timeline of the recording schedule. Mice were entrained to opposing 12 h:12 h light–dark cycles such that recordings could be obtained across the circadian cycle. Three mice began recordings at CT6 and two at CT18 (black arrows). Stars indicate 10-min recording epochs. D. Representative spike rasters (top) and PSTHs (bottom) from two electrode channels from two mice showing responses to light (0.5s) during electrode implantation. E. Mean firing rate of all light-responsive dLGN recording channels from each mouse during successive 10-min recording epochs across 24 h. Black line indicates the population mean; pink line shows the fitted cosinor model. Rhythm detection: *F* = 29.2, *p* = 0.0108; rhythm percentage: *r* = 0.86, PR = 0.74, *p* = 3.31 × 10⁻⁴. F. Representative trajectories of locomotor activity from two mice during the subjective day (left) and subjective night (right) during 10-min recording epochs. G. Mean z-scored distance travelled during each 10-min recording epoch. Locomotor activity exhibited a circadian rhythm, peaking during the subjective night. Black line indicates the population mean; pink line shows the fitted cosinor model. Rhythm detection: *F* = 17.6, *p* = 0.022; rhythm percentage: *r* = 0.839, PR = 0.703, *p* = 6.54 × 10⁻⁴. PR, percent rhythm; CT, circadian time; FR, firing rate.

Histological verification confirmed recordings from 48 light-responsive channels distributed throughout the dLGN in five mice (Fig 1D, Fig S1A). Notably, the firing rate of these light-responsive channels exhibited significant circadian variation across the 24-hour cycle (Fig 1E; cosinor analysis, rhythm detection P = 0.0108), with peak activity occurring in the late subjective day (CT10.8), demonstrating a robust circadian rhythm in spontaneous dLGN activity.

Locomotor activity is known to modulate dLGN activity (Aydın et al., 2018; Liang et al., 2020; Orlowska-Feuer et al., 2022; Reinhold et al., 2023), and is itself strongly regulated by circadian time, hence we next asked the extent to which locomotor vs. time-of-day effects might each define spontaneous firing. As expected, mice displayed robust circadian rhythms in locomotor activity, assessed both during the 10-minute recording epochs (Fig 1F,G; cosinor analysis, rhythm detection P = 0.02) across the intervening inter-epoch periods (Fig S1B, cosinor analysis, P = 0.025). Importantly, however, the peak of locomotor activity occurred around 9 hours after the peak in spontaneous firing activity, suggesting the two rhythms were not tightly coupled. We further found that mice spent significantly more time inactive during the subjective day than the subjective night (Fig S1C,D, paired t-test, p = 0.0025), indicating that circadian behavioural organisation remained intact throughout the experimental protocol. Crucially, recording epochs with greater locomotor activity were not associated with increased neuronal firing rates, but in fact a slight decrease (Fig S1E; Pearson’s correlation, p = .013). Together, these data demonstrate that spontaneous activity in the dLGN exhibits a robust circadian rhythm that cannot be explained by concurrent locomotor state, establishing circadian phase as an independent determinant of baseline dLGN excitability.

### Elevated daytime dLGN neural activity is maintained across visual contexts

Having established a circadian rhythm in spontaneous dLGN activity in awake animals, we next asked whether circadian phase also alters visually evoked activity under controlled stimulus conditions. To achieve precise control over visual stimulation, we performed extracellular recordings from dLGN neurons in anaesthetized mice at either ZT6 or ZT18, corresponding respectively to the ascending and descending phases of the circadian rhythm observed in awake animals. Recordings were obtained from animals maintained under standard light-dark conditions with controlled light history prior to experimentation, yielding 367 well-isolated single units distributed throughout the dLGN (Fig 2A,B; ZT6: n=204 units, n=6 mice; ZT18: n= 163 units, n=6 mice). We examined visual responses across a broad range of conditions by presentation of a stimulus battery comprising steady light, white-noise, temporal chirp and contrast chirp stimuli superimposed upon a staircase of increasing and decreasing background irradiances spanning approximately moonlight to daylight intensities (Orlowska-Feuer et al., 2023) (Fig. 2C,D). This stimulus was designed to evoke diverse patterns of neuronal activity (Fig 2E) and assess visual processing across a range of behaviourally relevant visual contexts.

**Figure 2.**
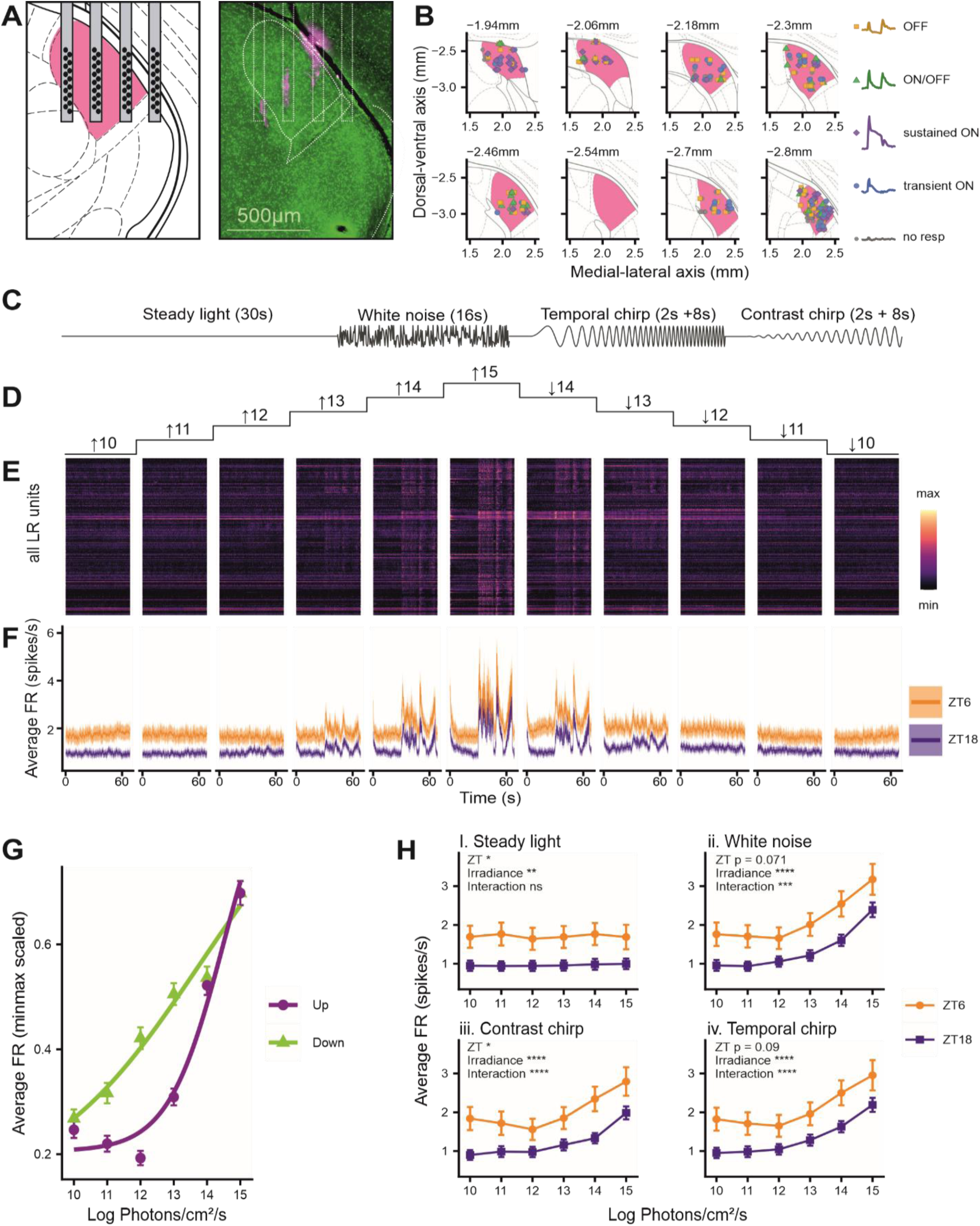
Elevated daytime firing of light-responsive dLGN neurons is maintained across visual stimuli and irradiance levels. Orange, ZT6 (midday); blue, ZT18 (midnight). A. Schematic showing electrode placement within the dLGN and the layout of recording contacts on a single electrode shank (left), together with a representative histological section showing DiI-labelled electrode tracts within the dLGN (right; DAPI, green; DiI, magenta). B. Projected anatomical locations of all isolated dLGN units classified according to light response type. Coordinates are given relative to bregma; panel headings indicate anterior-posterior position. Points are coloured according to response classification. C. Schematic showing the intensity changes occurring within the four visual stimuli presented at each background irradiance step. D. Background irradiance (log photons/cm^2^/s) during the ascending and descending phases of the irradiance staircase. E. Heatmaps of normalized PSTHs from all light-responsive dLGN units across the complete visual stimulus battery (C,D). For each unit, firing rates were normalized to the unit-specific minimum and maximum values; warmer colours indicate higher firing rates. F. PSTHs showing the population-average firing rate of light-responsive dLGN neurons recorded at midday (orange, ZT6, n = 136) or midnight (blue, ZT18, n = 120) across the staircase stimulus, revealing consistently elevated activity at midday. G. dLGN firing exhibits distinct irradiance-response relationships during the ascending (purple) and descending (green) phases of the staircase stimulus. Average firing rates were calculated across the complete stimulus battery at each irradiance step. Separate Hill fits were favoured over a shared model (F(4,3016) = 29.0, P = 1.14 × 10⁻²³). H. Circadian modulation of firing rate is evident across all classes of visual stimulus. Mean firing rates of light-responsive neurons are shown for each ascending irradiance step during (i) steady light (mixed ANOVA: ZT, F(1,250) = 6.2, P = 0.013; irradiance, F(2.13,532.85) = 5.9, P = 0.002; ZT × irradiance, F(2.13,532.85) = 1.3, P = 0.282), (ii) temporal white noise (mixed ANOVA: ZT, F(1,250) = 3.3, P = 0.072; irradiance, F(2.02,503.96) = 235.0, P = <0.001; ZT × irradiance, F(2.02,503.96) = 7.9, P <0.001) (iii) contrast chirp (mixed ANOVA: ZT, F(1,250) = 4.8, P = 0.029; irradiance, F(2.25,562.65) = 149.4, P = <0.001; ZT × irradiance, F(2.25,562.65) = 9.4, P <0.001;) and (iv) temporal frequency chirps (mixed ANOVA: ZT, F(1,250) = 2.9, P = 0.092; irradiance, F(2.09,523.64) = 187.7, P <0.001; ZT × irradiance, F(2.09,523.64) = 9.2, P <0.001). Statistical analyses were performed on log-transformed firing rates; data are displayed in the original scale.

Consistent with our observations in awake animals, light-responsive neurons recorded at ZT6 generally exhibited higher firing rates than those recorded at ZT18, irrespective of the visual stimulus presented or the prevailing background irradiance (Fig. 2F; Fig S2A,B,C,D). Interestingly, the relationship between irradiance and firing rate differed between ascending and descending irradiance transitions (Fig. 1G; nested model comparison, P = 8.39 × 10⁻²⁵), consistent with previous reports using similar stimuli (Orlowska-Feuer et al., 2023; Schmidt et al., 2014). Because the origin of this asymmetry remains unclear and was not influenced by circadian phase, subsequent analyses focused exclusively on the ascending irradiance series.

Diurnal modulation of firing was evident across all visual contexts examined. During steady-light stimulation, neurons recorded at ZT6 exhibited elevated firing rates across irradiance levels, resulting in a significant main effect of circadian phase without evidence for a ZT × irradiance interaction (Fig. 2H.i). For white-noise (Fig 2H.ii), contrast chirp (Fig2H.iii) and temporal chirp stimuli (Fig 2H.iv), the magnitude of the day-night difference varied across irradiance levels, producing significant ZT × irradiance interactions. Nevertheless, across these stimulus classes, firing rates remained generally higher in neurons recorded during the day than at night. Together, these findings demonstrate that circadian phase is an important determinant of dLGN activity across a broad range of visual stimuli and ambient light levels, and this modulation can operate independently of irradiance-dependent changes in firing.

### Polarity, contrast and temporal tuning are preserved across circadian phase

We next asked whether circadian phase altered the fundamental visual response properties of dLGN neurons, or whether its effects were restricted to baseline activity. Using responses to light steps from darkness, we found that the proportions of different response types were comparable between ZT6 and ZT18 (Fig. 3A, 3B; Chi-square, p = 0.26). Likewise, the distribution of response polarity angles was similar across timepoints (fig 3C; p > 0.10), with units clustered predominantly within the ON+ region indicating that excitatory responses to light onset represented the dominant response profile. The prevalence of each response class was also consistent with previous reports from the dLGN (Grubb and Thompson, 2003; Piscopo et al., 2013), indicating that circadian phase altered baseline activity without changing the overall composition of visual response types.

**Figure 3.**
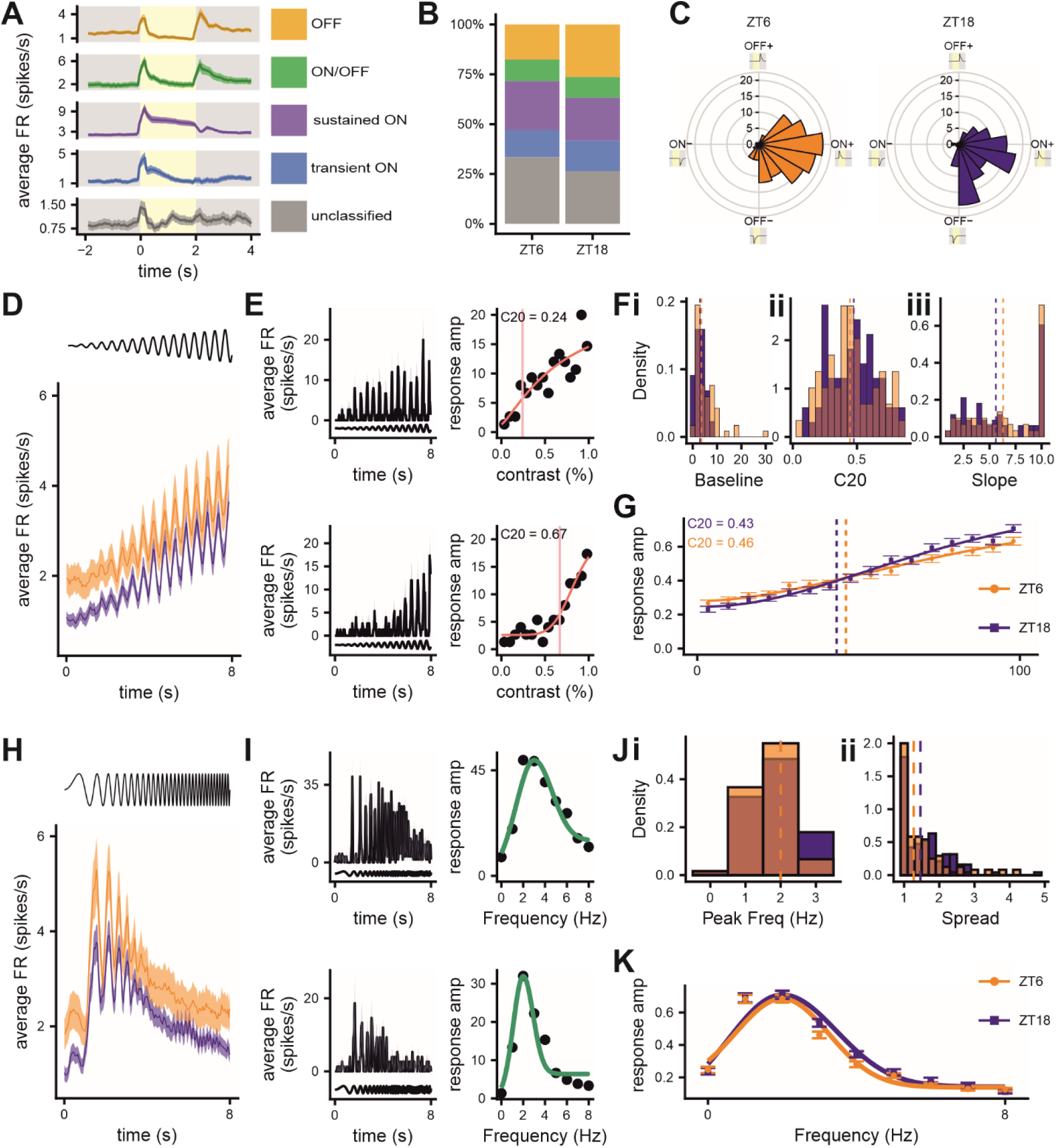
Core visual response properties of dLGN neurons are preserved across circadian phase despite changes in baseline activity. Orange, ZT6 (midday); blue, ZT18 (midnight). A. Average PSTHs of all dLGN neurons classified by light response type. B. The proportions of each light response type recorded in the dLGN were not significantly altered by the time of day (ZT6, n = 204; ZT18 =163; Chi-square: χ²(4) = 5.19, *P* = 0.268). C. Response polarity angle distributions did not differ between circadian phases (Watson two-sample test, *U*² = 0.082, *P* > 0.10) D. PSTHs of all light-responsive dLGN neurons recorded either at midday or midnight in response to the contrast chirp stimuli at the brightest irradiance (15 log photons/cm^2^/s). Timing of the contrast chirp stimulus shown above panel. E. Representative PSTHs of two dLGN neurons at the brightest contrast chirp stimulus (left) and corresponding contrast sensitivity functions (right). Pink line marks the estimated C20 value. F. Contrast sensitivity function parameters were estimated for individual neurons with model fits of R² > 0.5 (ZT6, n = 59 ZT18, n = 66). Baseline amplitudes (i) differed between circadian phases (Wilcoxon rank-sum test, *W* = 1429, *P* = 0.011), whereas C20 (ii) and slope (iii) did not (*W* = 2044, *P* = 0.63; *W* = 1746, *P* = 0.31, respectively). Dotted lines indicate group medians. G. Population contrast sensitivity functions fitted to normalized responses of all light-responsive dLGN neurons recorded at ZT6 or ZT18. Dotted lines indicate the estimated C20 values. H. PSTHs of all light-responsive dLGN neurons in response to temporal frequency chirp stimuli at the brightest irradiance. Timing of the temporal chirp stimulus shown above panel. I. Representative PSTHs of two dLGN neurons at the brightest temporal chirp for two individual dLGN units (left) and their respective temporal tuning curves (right). J. Frequency tuning parameters estimated for individual neurons with R² > 0.5 (ZT6, n = 120; ZT18, n = 95). Preferred temporal frequency (i) and tuning spread (ii) did not differ between timepoints (Wilcoxon rank-sum test, preferred frequency: *W* = 6351, *P* = 0.11; spread: *W* = 6013, *P* = 0.48). Dotted lines indicate group medians. K. Population temporal frequency tuning curves fitted to normalized responses of all light-responsive dLGN neurons recorded at ZT6 and ZT18.

We next quantified contrast-response functions using responses to the contrast chirp stimulus (Fig 3D, E). At the individual-unit level, whilst baseline response amplitudes differed substantially between timepoints (Fig. 3F.i), the other fitted parameters (slope and C20) remained largely unchanged (Fig. 3F.ii-iii). Consistent with this, population-average contrast-response functions of all dLGN light-responsive units yielded similar C20 estimates, with 20% of the maximal response reached at 43% contrast at ZT18 and 46% contrast at ZT6. These findings thus provide little evidence for circadian differences in contrast sensitivity or the overall shape of the contrast-response relationship.

A similar pattern was observed for temporal frequency tuning (Fig 3H). Although individual neurons exhibited the expected diversity in tuning profiles, ranging from narrowly tuned to broadly responsive units (Fig. 3I), neither the preferred temporal frequency (Fig 3J.i. median peak frequency = 2 Hz at both timepoints; P = 0.11) nor tuning width (Fig 3Jii. ZT6 median = 1.27 Hz; ZT18 median = 1.47 Hz; P = 0.48) differed between timepoints. Population-level analyses yielded the same conclusion, revealing nearly identical temporal frequency tuning from dLGN light-responsive units at ZT6 and ZT18, with no apparent shift in either peak frequency or tuning width (Fig 3K).

Together, these findings indicate that circadian phase exerts a robust influence on overall dLGN activity while leaving fundamental contrast and temporal tuning properties largely unchanged.

### Adjustments in spatial acuity across circadian phase

Having found that contrast and temporal tuning were preserved across circadian phase, we next asked whether spatial integration was similarly stable. We applied a binary white noise spatial stimulus to map the linear receptive field of neurons at midday and midnight. Spike triggered averages revealed the receptive field size and dynamics for a sub-population of light-responsive units (Fig 4A-C; midday: n = 43 units, n= 5 mice; midnight: n = 50 units, n= 5 mice). Quantification of receptive field size revealed a significant expansion of spatial receptive fields at midnight relative to midday (median diameter = 8.09° vs 6.05°, respectively), indicating that dLGN neurons integrate visual information over a larger region of visual space during the night.

**Figure 4.**
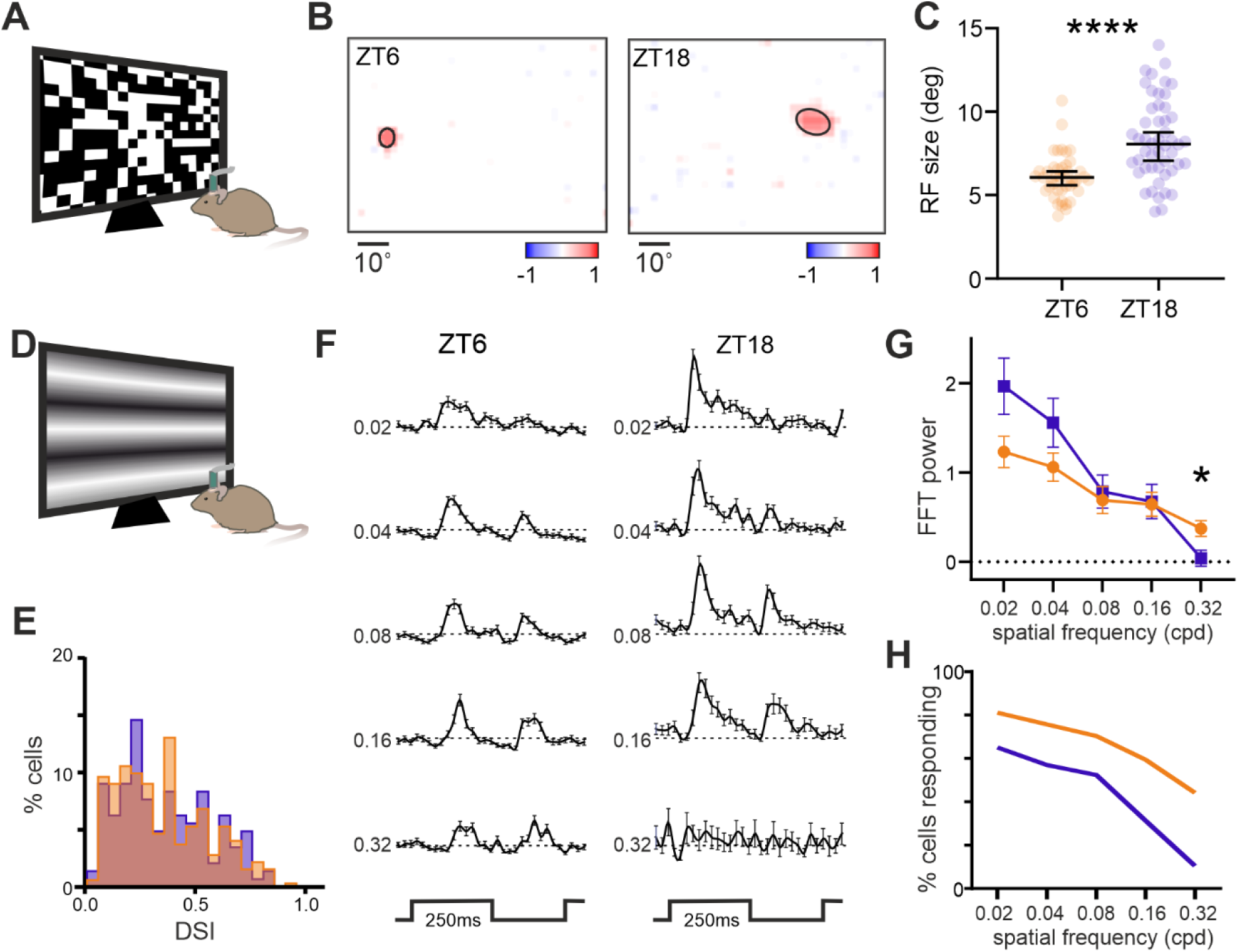
Circadian phase alters receptive field size and representation of fine spatial detail. Orange, ZT6 (midday); blue, ZT18 (midnight). A) Schematic depicting the binary white noise stimulus used to map receptive fields in the mouse dLGN. B) Representative spike-triggered averages recorded at ZT6 and ZT18. Black ellipses show two-dimensional Gaussian fits used to estimate receptive field size. C) Receptive fields were significantly larger at midnight than midday, indicating expanded spatial integration during the nocturnal phase (Mann-Whitney test, *P* < 0.001; ZT6, *n* = 43 units; ZT18, *n* = 50 units). D) Schematic illustrating the presentation of 2 Hz contrast-inverting sinusoidal gratings used to assess spatial frequency tuning. E) Orientation selectivity was preserved across circadian phase. Distribution of orientation selectivity indices for dLGN neurons recorded at ZT6 (*n* = 111) and ZT18 (*n* = 86) (Mann-Whitney test, *P* = 0.24). F). Mean ± SEM PSTHs evoked by 2 Hz inverting gratings across five spatial frequencies (0.02, 0.04, 0.08, 0.16 and 0.32 cpd) for neurons recorded at ZT6 and ZT18. Data are shown for each unit’s preferred stimulus orientation and phase. G) Responses to high spatial frequency stimuli were selectively enhanced during at midday. FFT amplitude of the F1 response component at the preferred stimulus orientation and phase is shown as a function of spatial frequency. A significant increase in response amplitude was observed at 0.32 cpd at ZT6 (mixed ANOVA: ZT, F(1,195) = 1.34, *P* = 0.25; spatial frequency, F(3.365,656.1) = 28.4, *P* < 0.001; ZT × spatial frequency, F(4,780) = 4.065, *P* = 0.0029; Sidak post hoc test, *P* = 0.04 at 0.32 cpd). H) Percentage of neurons responding at each spatial frequency tested at ZT6 vs. ZT18.

We next asked whether this change in receptive field size was accompanied by alterations in the encoding of spatially structured visual stimuli. Inverting sinusoidal grating stimuli were presented to mice at 2Hz, at a range of spatial frequencies and orientations (Fig 4D,F). We found that units in the dLGN had equivalent sensitivity range as previously described, with neurons tracking inverting stimuli across the full range tested. Units also demonstrated an equivalent degree of orientation selectivity at ZT6 and ZT18 (Fig 4E). However, we found that the response amplitude and number of units responding to the highest spatial frequency tested (0.32cpd) was significantly reduced at midnight (Fig 4G,H). Thus, the expansion of receptive fields observed at midnight was accompanied by a reduced representation of fine spatial detail.

### Augmented dynamic range in the circadian day

Changes in ongoing activity in the dLGN are not expected to alter contrast or temporal tuning *per se,* but they do generate predictions for how visual signals are encoded. One possibility is that elevated baseline firing can move neurons into a more linear operating regime, thereby extending their effective dynamic range. We tested this hypothesis by exploring the irradiance response relationship for light steps from dark, presenting light steps spanning six orders of magnitude in intensity.

We used responses at the brightest light step to identify light-responsive units and confirm recordings were drawn from similar populations of neurons at each timepoint. Although response class and polarity distributions differed significantly between phases (Fig 5A, B, Chi-square, p = 8.7 × 10⁻⁴; 5C, Watson two-sample test, P < 0.001), the overall composition of the recorded population was broadly similar, with sustained ON and transient ON cells remaining the predominant response classes at both ZT6 and ZT18.

**Figure 5.**
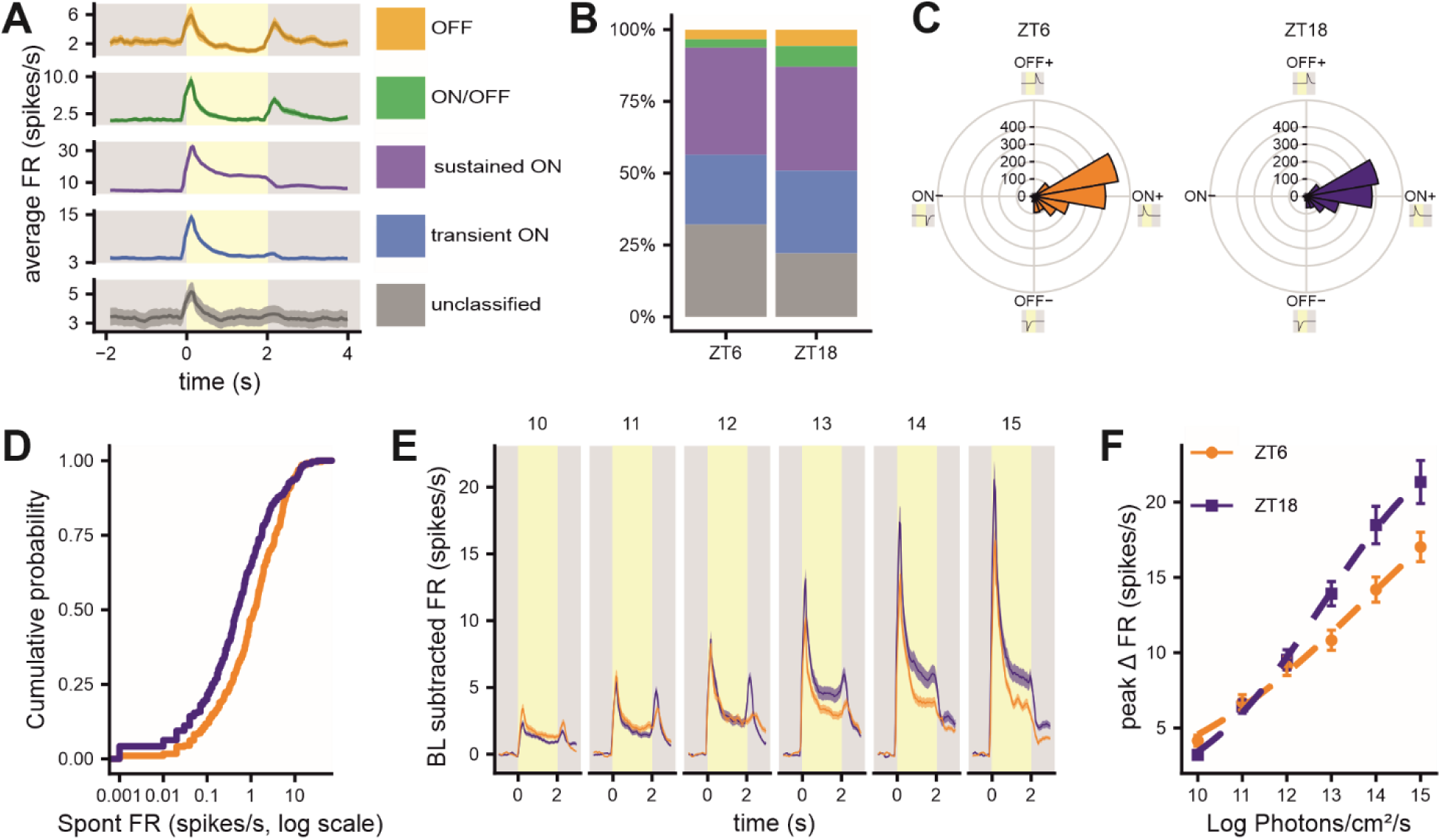
Dynamic range of light-responsive dLGN neurons is enhanced during the subjective day. Orange, ZT6 (midday); blue, ZT18 (midnight). A. Average PSTHs of all dLGN neurons grouped according to light response type. B. The distribution of light-response types recorded at each time point (ZT6, n = 411; ZT18, n = 385; Chi-square: χ²(4) = 18.77, P = 8.7 × 10⁻⁴). C. The distribution of response polarity angles differed between time points (Watson two-sample test, U² = 0.418, P < 0.001), although neurons remained predominantly clustered within the ON+ quadrant. D. Empirical cumulative distribution functions of spontaneous firing rates recorded in darkness from light-responsive dLGN neurons at ZT6 (n = 279) and ZT18 (n = 300). Firing rates are plotted on a log10 scale. Spontaneous firing rates were higher at ZT6 than ZT18 (Wilcoxon rank-sum test, W = 51,635, P = 5.6 × 10⁻⁷). E. Average baseline-subtracted PSTHs of light-responsive dLGN neurons evoked by a 2-s light step from darkness at ZT6 and ZT18. Irradiance (log photons/cm^2^/s) indicated above each panel. F. Peak firing rate during the first 500ms following light onset is plotted as a function of irradiance. Responses recorded at night saturated at lower irradiances, whereas neurons recorded during the day maintained graded increases in firing across a broader range of intensities, consistent with an expanded dynamic range (extra sum-of-squares *F*-test: F(3,3468) = 11.76, *P* = 1.16 × 10⁻⁷).

We once again found evidence for diurnal modulation of baseline excitability in this dataset. As spontaneous dark-adapted firing rates differed across circadian phase (Fig 5D), responses were quantified relative to each unit’s baseline activity, allowing direct comparison of stimulus-evoked response amplitudes between timepoints (Fig. 5E). For each light-responsive unit, we measured the peak increase in firing rate following light onset and fitted irradiance-response functions to the population data (Fig. 5F).

The relationship between irradiance and response amplitude differed significantly across circadian phase (P < 0.001). Responses recorded at night saturated at lower irradiances, whereas neurons recorded during the day maintained graded increases in firing across a broader range of light intensities, consistent with a shift in operating point associated with elevated baseline firing, placing neurons in a more linear response regime.

### Information transfer is reconfigured across the circadian cycle

The observed increases in baseline activity, together with selective changes in visual coding, suggested that circadian phase could influence both the amount and efficiency of information transmitted by dLGN neurons. We therefore asked how information transfer varied across the circadian cycle under different visual stimulus conditions.

We first analysed the temporally modulated full-field white-noise stimulus presented as part of our staircase stimuli (Fig 6A). This allowed precise measurement of information transfer rate under controlled statistical conditions (see methods). Consistent with our previous analyses, firing rates were higher during the day and increased with background irradiance (Fig 6B).

**Figure 6.**
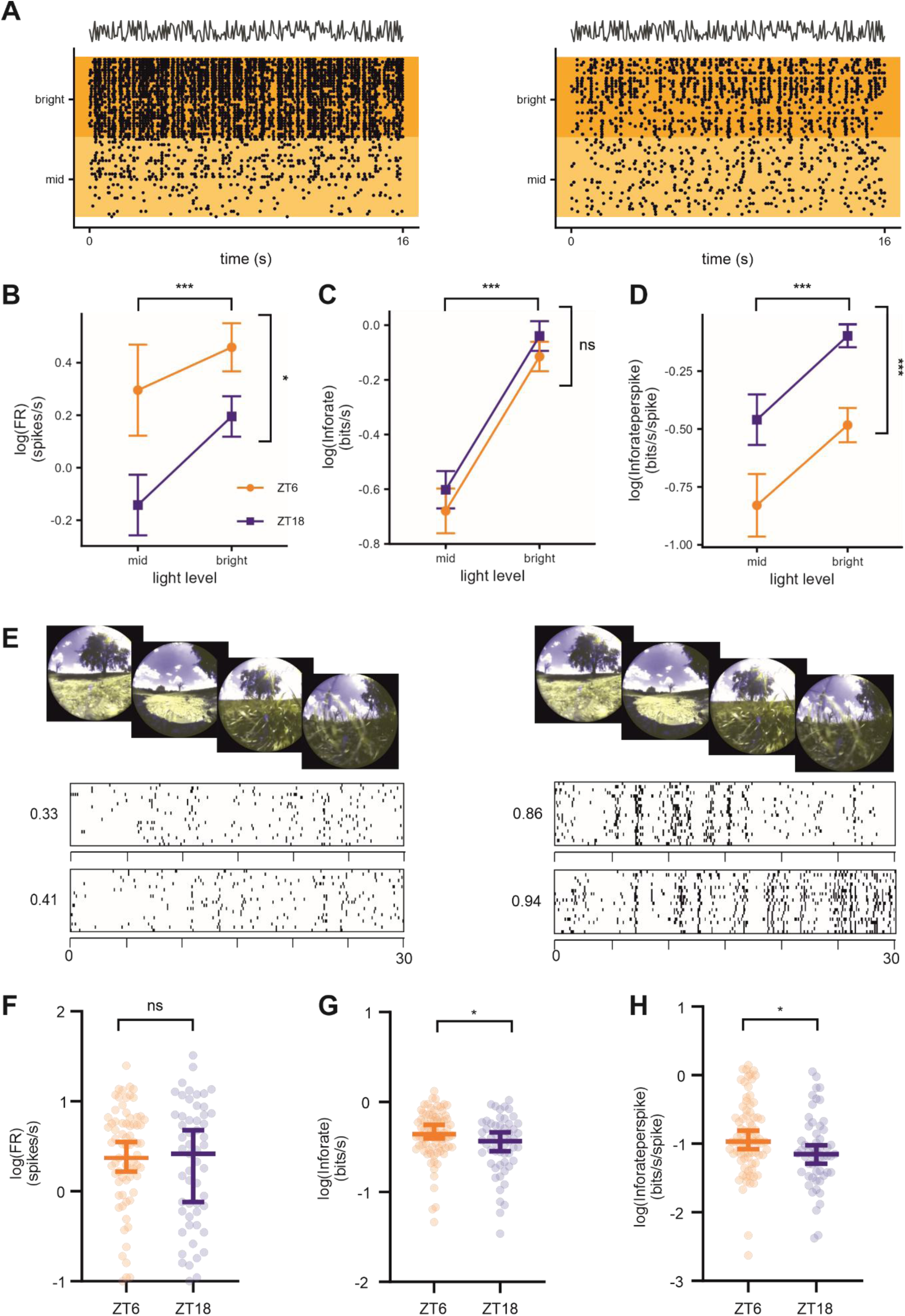
Circadian phase reconfigures information transfer in a stimulus-dependent manner. Orange, ZT6 (midday); blue, ZT18 (midnight). A. Representative spike rasters showing how two units responded to repeated presentations of the white noise stimuli either at “mid” (12 and 13 log photons/cm^2^/s; light yellow) or “bright” irradiance (log photons/cm^2^/s; dark yellow). Profile of white noise intensity changes shown above. B. Firing rates increased with irradiance and were consistently higher at midday than midnight (ZT6, *n* = 136; ZT18, *n* = 115; linear mixed model, irradiance: F(1,102.3) = 88.66, *P* < 0.001; circadian phase: F(1,251.8) = 4.42, *P* = 0.037; interaction: F(1,102.3) = 0.48, *P* = 0.489). C. Total information transfer increased with irradiance but was unaffected by circadian phase (linear mixed model, irradiance: F(1,151.9) = 115.33, *P* < 0.001; circadian phase: F(1,246.6) = 1.48, *P* = 0.224; interaction: F(1,151.9) = 0.04, *P* = 0.840). D. Coding efficiency (information transmitted per spike) increased with irradiance and was significantly greater at midnight, indicating more efficient information transfer rate under low-firing conditions (linear mixed model, irradiance: F(1,194.1) = 16.16, *P* = 8.34 × 10⁻⁵; circadian phase: F(1,223.3) = 17.00, *P* = 5.28 × 10⁻⁵; interaction: F(1,194.1) = 0.01, *P* = 0.933). E. Representative spike rasters showing responses of four dLGN neurons to repeated presentations of a naturalistic movie stimulus. Corresponding information transfer rates (bits s⁻¹) are shown to the left of each raster. Example movie frames are shown above (adapted from Qiu *et al*., 2021). F. Firing rates during presentation of the natural movie were similar at ZT6 (*n* = 79) and ZT18 (*n* = 55) (Mann-Whitney test, *P* = 0.11). G. In contrast to white-noise stimulation, information transfer during naturalistic stimulation was significantly greater at midday than midnight (Mann-Whitney test, *P* = 0.04). H. Coding efficiency (bits/spike) during naturalistic stimulation was also significantly enhanced at midday (Mann-Whitney test, *P* = 0.02), indicating that the effects of circadian phase on information coding depend upon stimulus statistics.

Despite these differences in firing rate, total information transmitted (bits/s) was unaffected by circadian phase (Fig 6C), indicating that increased daytime firing did not simply translate into greater information output. Instead, coding efficiency (bits/spike) was significantly higher at night (Fig 6D), showing that individual spikes carried more information under low-firing conditions in response to this stimulus. By comparison, irrespective of circadian time, increasing background irradiance increased total information transmitted, consistent with previous observations in the retina (Milosavljevic et al., 2018).

Thus, under the relatively simple statistical structure of full-field white noise, circadian phase altered coding efficiency without changing overall information transfer rate. We next asked whether this relationship was maintained for visual inputs with richer spatial and temporal structure by presenting a naturalistic movie (Fig 6E) approximating the visual experience of a mouse in a daytime environment (Qiu et al., 2021). Restricting our analysis to neurons which responded to binary white noise/inverting grating stimuli (Fig 4), we first confirmed that this subset showed the same stable information transfer rate across circadian phase for temporal white-noise stimuli (Linear mixed effects model; Šídák’s multiple comparisons test; p= 0.86 ZT6: n=79 units; ZT18: n=55 units). Overall information transfer rate during the natural movie was lower than during white-noise stimulation (mean = 1.2 vs 0.4 bits/s; Linear mixed effects model; effect of stimulus type = p<0.001), consistent with its lower contrast and greater spatial complexity. Importantly, however, the natural movie revealed a circadian effect that was absent for white noise: neurons recorded during the day transmitted significantly more information than those recorded at night, accompanied by a corresponding increase in the rate of information transfer. Thus, while circadian phase primarily alters coding efficiency under simple stimulus conditions, it enhances overall information transfer rate when neurons encode more naturalistic visual scenes.

These findings therefore indicate that the impact of circadian phase on information transfer depends on stimulus statistics. Under white-noise stimulation, circadian modulation primarily affects coding efficiency, whereas during naturalistic stimulation it increases overall information transfer rate, indicating that the influence of circadian state on visual coding is strongly context dependent.

### Circadian modulation is expressed at the retinogeniculate synapse

The circadian changes in visual coding we observed could reflect alterations in retinal output, intrinsic thalamic properties, or transmission at the retinogeniculate synapse. We therefore asked whether circadian modulation is evident at the retinogeniculate synapse itself by selectively activating retinal ganglion cell (RGC) terminals optogenetically while recording from dLGN neurons. Extracellular recordings were performed in Brn3c-Cre × ReaChR mice, which express the light-sensitive opsin ReaChR in a subset of RGCs (Fig. 7A; (Parmhans et al., 2020)). Using a fibre-coupled recording probe (“optrode”), we delivered light pulses directly within the dLGN while simultaneously recording neuronal activity (Fig. 7B). By directly activating RGC terminals within the dLGN, this approach bypasses retinal visual processing while allowing transmission across the retinogeniculate synapse to be assessed directly.

**Figure 7.**
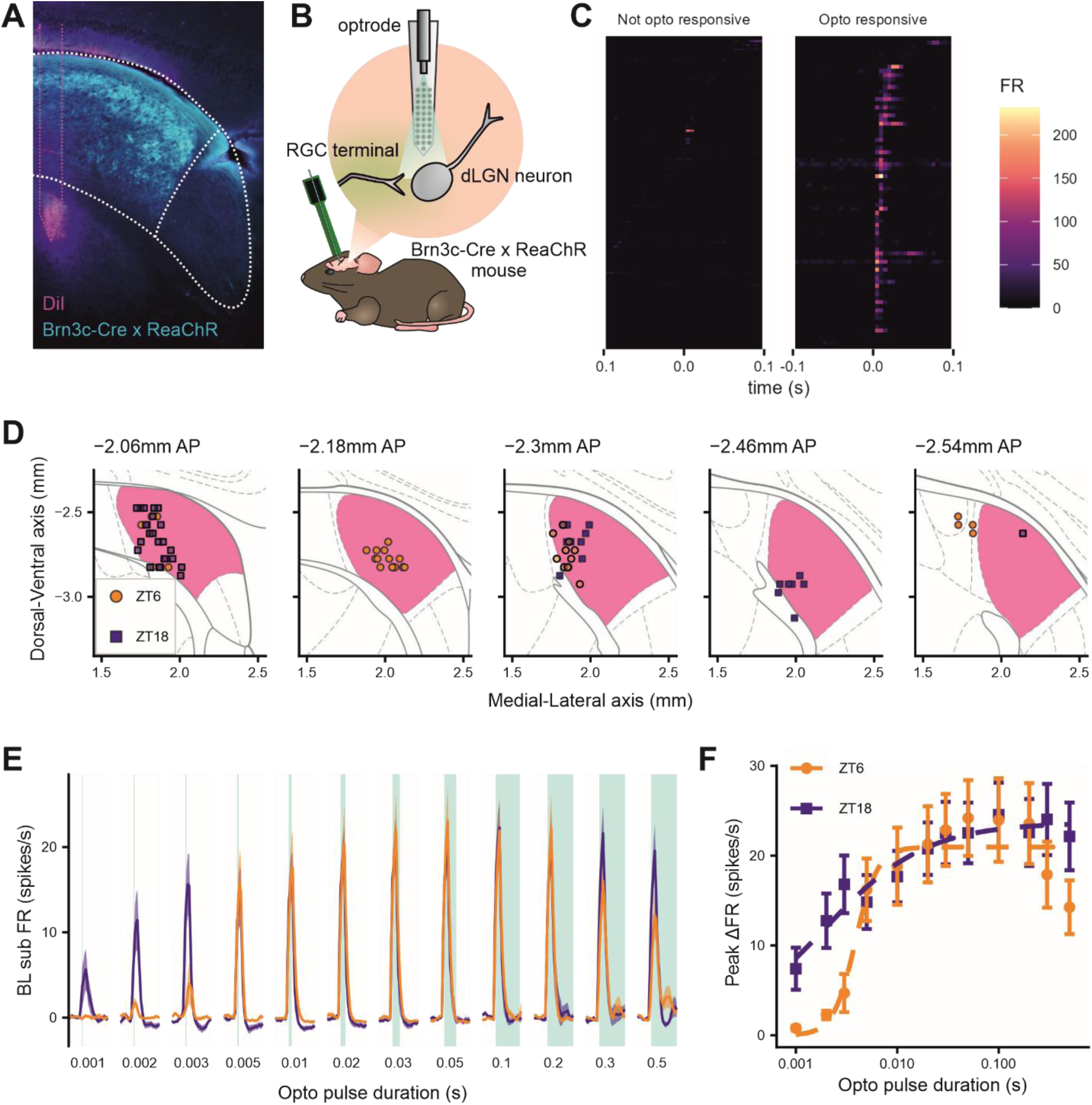
Circadian modulation of visual coding is expressed at the retinogeniculate synapse. Orange, ZT6 (midday); blue, ZT18 (midnight). A Representative histological image showing electrode tracts (magenta) and Brn3c^Cre^-positive retinal ganglion cell terminals within the dLGN visualised by mCitrine fluorescence (cyan). B. Schematic illustrating how optogenetic activation of Brn3c^Cre^-ReaChR retinal ganglion cell terminals within the dLGN was combined with extracellular recordings to assess retinogeniculate transmission. C. Heatmaps showing average PSTHs of dLGN neurons in response to 50 presentations of a 200ms optogenetic pulse. Units were classified as non-responsive (left) or opto-responsive (right). D. Projected anatomical locations of all opto-responsive dLGN units recorded at ZT6 and ZT18. Coordinates are given relative to bregma; panel headings indicate anterior-posterior position. E. Population mean response PSTHs of opto-responsive dLGN neurons recorded at midday (ZT6, n = 35) or midnight (ZT18, n = 42) in response to optogenetic pulses of increasing duration. F. Circadian phase altered the optogenetic input-output relationship of dLGN neurons. Peak firing rate responses are plotted as a function of optogenetic pulse duration. Neurons recorded at ZT18 exhibited larger responses to brief optogenetic stimuli than those recorded at ZT6, consistent with enhanced retinogeniculate gain during the nocturnal phase (extra sum-of-squares F-test, F(3,842) = 3.18, P = 0.023).

Units were recorded throughout the dLGN (Fig 7D, Fig S3D) and exhibited diurnal modulation of visually evoked responses comparable to those we observed in wild-type animals (Fig S3A,B,C,E,F). Futhermore, expression of ReaChR did not measurably alter visually evoked response properties relative to wild-type controls (Fig. S3G).

We identified neurons that responded directly to optogenetic stimulation (Fig. 7C,D; see Methods) and quantified the peak firing rate evoked by light pulses of increasing duration (Fig. 7E). Circadian phase significantly altered the optogenetic input-output relationship (extra sum-of-squares F-test: *p* = 0.023), with neurons recorded at night exhibiting larger responses to brief stimuli than those recorded during the day (Fig. 7F). Thus, equivalent inputs elicited stronger postsynaptic responses at night, indicating that transmission across the retinogeniculate pathway is enhanced during the nocturnal phase, consistent with increased synaptic gain at the retinogeniculate synapse.

## Discussion

Neuronal firing and functional connectivity vary dynamically with changes in brain state. Here, we identify the circadian clock as a novel locus of control that drives anticipatory daily changes in the function of the mouse primary visual thalamus (dLGN). Using *in vivo* recordings from awake mice maintained in constant darkness across 24 hours, we show that dLGN neurons exhibit robust circadian rhythms in spontaneous firing that peak just before the subjective day-night transition. Although this modulation persists during ongoing visual stimulation, it does not globally reconfigure visual tuning: contrast sensitivity, temporal frequency tuning, and response classifications remain remarkably stable across circadian phase. Instead, circadian time selectively alters other dimensions of visual coding, including receptive field size, spatial frequency representation, response dynamic range, and the efficiency with which visual information is transmitted. In particular, nighttime responses are characterised by larger receptive fields, enhanced information efficiency, and increased sensitivity to weak retinogeniculate inputs, whereas daytime responses exhibit higher firing rates, broader dynamic range, and improved representation of fine spatial detail. Together, these findings demonstrate that circadian phase actively reconfigures the operating state of the retinogeniculate pathway, reshaping how visual information is represented and transmitted across the day-night cycle.

### Circadian control of spontaneous neural firing in the dLGN

Recent work has highlighted the ubiquity of circadian rhythms in neural activity across the brain (Begemann et al., 2020; Guilding and Piggins, 2007; Mendoza, 2025; Paul et al., 2020; Yamashita et al., 2025). Consistent with this body of work, we find that spontaneous firing in the dLGN exhibits a robust circadian rhythm, peaking during the late subjective day in both constant darkness and during ongoing visual stimulation. These findings extend previous evidence for rhythmicity within the visual thalamus. Chrobok and colleagues demonstrated rhythmic clock gene expression within the LGN under constant conditions, although the loss of coherent electrical rhythms in isolated dLGN slices suggested that this region may not contain a fully autonomous circadian oscillator (Chrobok et al., 2021). Instead, the dLGN may function as a downstream or “slave” oscillator, integrating timing information from other circadian centres, as has been proposed for several extra-SCN brain regions (Guilding and Piggins, 2007; Paul et al., 2020). Our findings are also broadly consistent with earlier electrophysiological work demonstrating diurnal variation in LGN firing activity in anaesthetized mice (Brown et al., 2011), as well as recent whole-brain c-Fos mapping studies identifying the dLGN as a rhythmic visual structure (Yamashita et al., 2025).

Importantly, the rhythmic modulation in dLGN activity we observed could not be explained by concurrent behavioural state. Neural activity showed only a weak relationship with locomotor behaviour, despite previous reports demonstrating movement-dependent modulation of dLGN activity (Crombie et al., 2024; Erisken et al., 2014; Niell and Stryker, 2010; Roth et al., 2016; Williamson et al., 2015), likely because our recordings sampled a heterogeneous population of neurons with diverse responses to locomotion. Furthermore, these time-of-day differences persisted under anaesthesia, indicating that the observed rhythm is unlikely to arise solely from previously described effects of movement, behavioural state or arousal (Bezdudnaya et al., 2006; Crombie et al., 2024; Durand et al., 2016; Liang et al., 2020; Molnár et al., 2021; Nestvogel and McCormick, 2022; Schröder et al., 2020). Rather, these data support the idea that circadian phase acts as an independent modulator of baseline dLGN excitability, providing an anticipatory signal that shapes thalamic processing across the day-night cycle.

### Coding efficiency is reorganised across the circadian cycle

A key question arising from the elevated daytime firing rates was whether they increase the amount of visual information transmitted through the dLGN. In many sensory systems, higher firing rates are associated with increased information transfer, albeit at substantial metabolic cost (Harris et al., 2015; Laughlin et al., 1998; Levy and Baxter, 1996; Milosavljevic et al., 2018). Our data, however, reveal that this relationship depends critically on the statistical structure of the visual stimulus. Under a high-contrast white-noise stimulus, neurons conveyed comparable amounts of information during day and night. Given the reduced firing at night, this resulted in greater efficiency in information transfer, with more information conveyed per spike. Thus, under simple stimulus statistics, circadian modulation primarily redistributes coding efficiency rather than increasing coding capacity.

In contrast, when neurons were driven by a naturalistic movie stimulus, information transfer was greater during the day. This suggests that circadian modulation becomes functionally significant when neurons are required to encode stimuli with richer spatial and temporal structure. Under these conditions, elevated firing rates, broader dynamic range, and enhanced representation of fine spatial detail may together support the encoding of more complex visual inputs.

These findings indicate that circadian modulation of information transfer is not fixed, but is revealed in a stimulus-dependent manner. Rather than simply scaling information in proportion to firing rate, the visual thalamus adjusts how information is distributed according to both internal state and the statistical structure of the visual input. Such flexibility may allow the system to balance coding efficiency against representational capacity in a manner that is matched to the demands of the visual environment (Attwell and Laughlin, 2001; Harris et al., 2015; Laughlin et al., 1998).

### The visual code is selectively reconfigured across the circadian cycle

A striking feature of the circadian effects described here was their selectivity. Despite robust changes in spontaneous firing rate and information transfer, several fundamental response properties remained remarkably stable across circadian phase. The proportions of ON, OFF and ON-OFF neurons were unchanged, as were measures of contrast sensitivity and temporal frequency tuning. These findings suggest that circadian modulation does not globally reorganise the functional architecture of the dLGN or alter the range of visual features represented within the population.

Instead, circadian phase selectively modified specific aspects of visual coding. Most notably, receptive fields were significantly larger at night, and this was accompanied by a reduction in representation of fine spatial detail. We additionally observed a broader intensity-response range during the day. Together, these findings indicate that circadian modulation acts primarily on the way visual information is represented rather than on the *identity* of the features encoded.

The dissociation between stable temporal and contrast tuning and altered spatial integration may provide important clues regarding the underlying mechanisms. Changes in receptive field size are likely to reflect alterations in the balance of convergence in retinal circuits or local inhibition, which are known to vary according to circadian phase. By contrast, the preservation of temporal tuning provides little evidence for a large-scale reconfiguration of retinal output or changes to intrinsic membrane properties of retinal neurons. Circadian signals therefore appear to selectively target those circuit elements that determine response gain and spatial pooling while leaving core feature selectivity largely intact.

Collectively, these findings suggest that circadian phase may alter the balance between visual sensitivity and spatial resolution within the retinogeniculate pathway. Larger receptive fields, enhanced coding efficiency and lower activation thresholds at night are broadly consistent with increased spatial pooling, a strategy that can improve reliability under noisy conditions at the expense of spatial precision. Conversely, daytime responses were characterised by broader dynamic range and enhanced representation of fine spatial detail. Although the behavioural consequences remain unknown, the coordinated nature of these effects suggests that circadian modulation acts on the visual system in a functionally meaningful manner rather than simply scaling neuronal excitability.

### Origins of circadian rhythms in the dLGN

The selective changes in visual coding described here raise the question of where within the visual pathway circadian modulation is implemented. Alterations in retinal output, local thalamic circuitry, neuromodulatory input, or synaptic transmission could all contribute to the observed changes. Our optogenetic experiments provide evidence that at least part of this modulation is expressed at the retinogeniculate synapse itself. By directly activating retinal ganglion cell terminals within the dLGN, we were able to bypass visual processing in the retina and assess the transformation of retinal input into thalamic output. Under these conditions, dLGN responses exhibited a clear circadian difference in their input-output relationship, with neurons recorded at night displaying lower activation thresholds and enhanced responses to weak optogenetic stimulation. These findings indicate that the efficacy of retinogeniculate transmission varies across circadian phase and identify the retinogeniculate synapse as a previously unrecognised locus of circadian regulation within the visual system.

Retinal neurons express autonomous molecular clocks and exhibit widespread daily rhythms in physiology, including changes in neurotransmitter release, photoreceptor function and retinal ganglion cell activity (Besharse and McMahon, 2016; Ruan et al., 2006; Storch et al., 2007). Such rhythms could also alter the strength or temporal structure of signals arriving at the dLGN and thereby contribute to the time-of-day-dependent changes observed here. The LGN itself expresses clock genes and exhibits circadian rhythms in molecular activity (Chrobok et al., 2021), raising the possibility that local clock-controlled processes influence synaptic efficacy, network excitability, or visual coding. In addition, the dLGN receives dense neuromodulatory input from arousal- and state-regulating centres, including cholinergic, serotonergic, orexinergic and noradrenergic pathways (Orlowska-Feuer et al., 2019; Pape and McCormick, 1989; Reggiani et al., 2023; Sokhadze et al., 2022), many of which exhibit pronounced daily variation in activity. These pathways are well positioned to integrate circadian and behavioural signals and shape thalamic processing accordingly.

## Conclusion

Together, our findings reveal circadian phase as a previously underappreciated determinant of visual thalamic function. Across the day-night cycle, circadian signals reshape spontaneous activity, dynamic range, spatial integration, information transfer and retinogeniculate gain, but leave many core aspects of visual tuning intact. Rather than globally altering visual feature selectivity, circadian modulation appears to selectively reconfigure how visual information is represented and transmitted within the dLGN. Together, these results suggest that the retinogeniculate pathway is a dynamic circuit whose operating state varies across the day-night cycle, and indicate that circadian time is a key determinant of sensory processing and circuit operating state.

## Acknowledgements

We would like to thank Dr Tudor Badea at Transylvania University of Brasov for the generous sharing of Brn3C^cre^ mice, and members of the BSF at the University of Manchester for assistance in husbandry and colony maintenance. The Bioimaging Facility microscopes used in this study were purchased with grants from BBSRC, Wellcome and the University of Manchester Strategic Fund, and we thank the Bioimaging Facility staff for their assistance with microscopy.

This work was supported by a Sir Henry Dale Fellowship 218556/Z/19/Z, jointly funded by the Wellcome Trust and the Royal Society (AEA).

## Author contributions

AEA supervised the project; AP and AEA designed the study and performed electrophysiological data collection and analysis, AP, JR, RS and AEA performed computational analysis of neural activity. AP and AEA wrote manuscript with input and approval of all authors.

---

**Figure S1.**
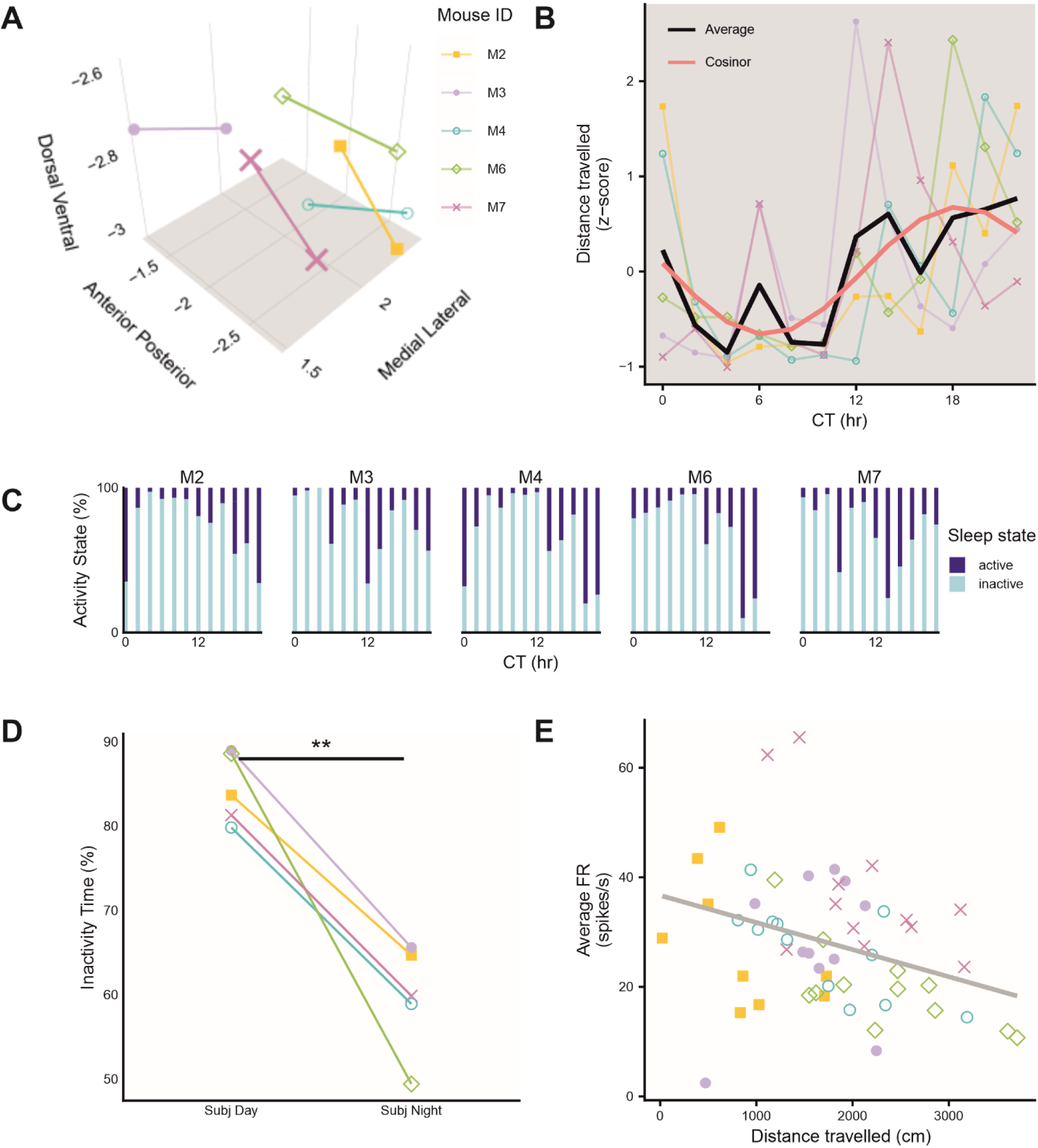
Electrode reconstruction and locomotor analyses for chronic recordings. A. Estimated rostral and caudal electrode tip locations reconstructed from histology. Lines indicate the inferred trajectories of the electrode arrays (based on the deepest point of each array) in 3D space. Coordinates are given relative to bregma. B. Mean distance travelled during the inter-epoch intervals. Mean z-scored locomotor activity exhibited a circadian rhythm, peaking during the subjective night. Black line indicates the population mean; pink line shows the fitted 24-h cosinor model. Rhythm detection: *F* = 16.0, *p* = 0.025; rhythm percentage: *r* = 0.812, PR = 0.660, *p* = 1.33 × 10⁻³. C. Each 10s of the inter-epoch intervals was classified as active or inactive according to the distance travelled. Figure shows the percentage each mouse spent in each activity state during the inter-epoch intervals. D. Mice spent a greater proportion of the inter-epoch intervals in the inactive state during the subjective day (CT0–CT12) than during the subjective night (CT12-24; paired two-sided *t*-test, *t*₄ = 6.76, *P* = 0.0025). E. Relationship between locomotor activity and spontaneous firing rate across 10-min recording epochs. Each point represents one recording epoch and plots the mean firing rate of light-responsive dLGN channels against the mean distance travelled during that epoch. Point shape identifies individual mice (see panel A). Grey line indicates the linear regression (Pearson correlation: *r* = −0.330, *P* = 0.013). CT, circadian time, PR, percent rhythm; FR, firing rate.

**Figure S2.**
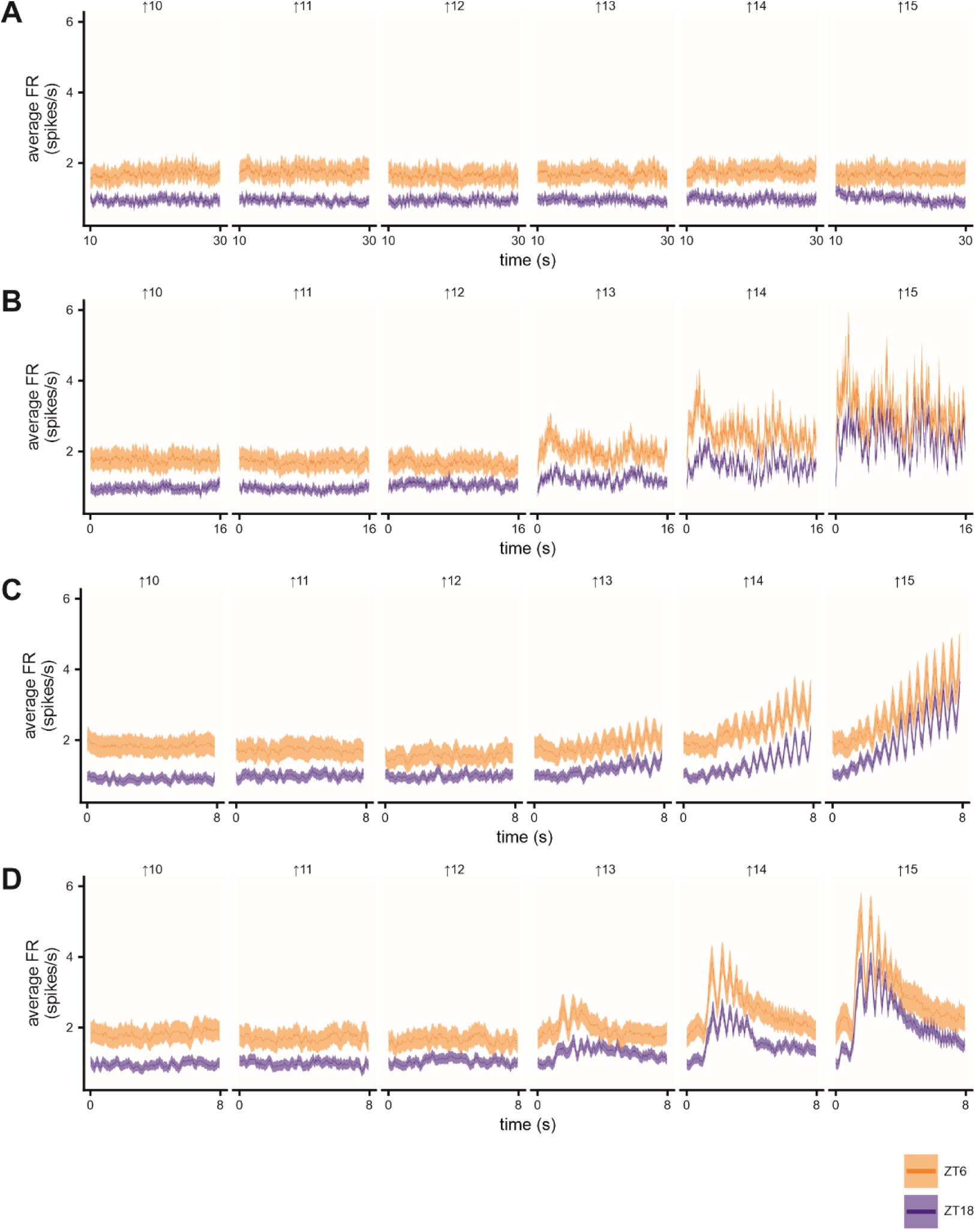
Population PSTHs of light-responsive dLGN neurons across visual stimuli and background irradiances. Population-average PSTHs of light-responsive dLGN neurons recorded during subjective day (orange, ZT6, n = 136) and subjective night (blue, ZT18, n = 120) across ascending background irradiances. Responses are shown for: (A) the final 20 s of steady illumination, (B) white-noise stimulation, (C) contrast chirp stimuli, and (D) temporal chirp stimuli.

**Figure S3.**
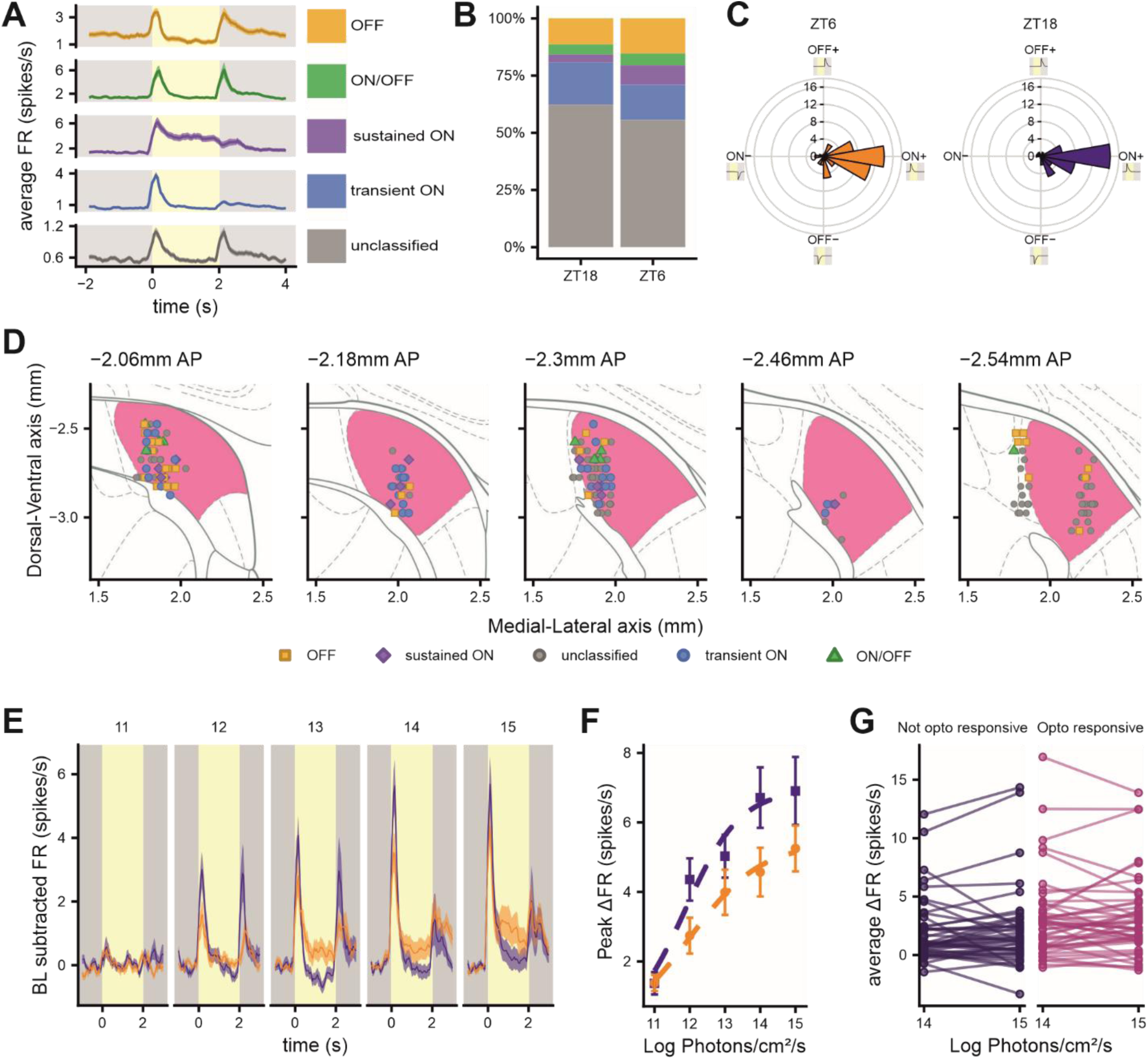
Brn3c^Cre^ x ReaChR mice had similar light responses to wildtype mice. Orange, ZT6 (midday); blue, ZT18 (midnight). A. Average PSTH of dLGN cells classified as each light response type. B. Similar proportions of light response types were observed at both timepoints (Chi-square: χ²(4) = 3.93, P = 0.416). C. Response polarity distributions were similar across times of day, with neurons predominantly clustered within the ON+ quadrant (Watson two-sample test, U² = 0.114, P > 0.10), in line with the dataset collected from the wildtype mice (Figure 5C). D. Projected anatomical locations of all dLGN units classified according to their light response. Coordinates are given relative to bregma, panel titles indicate location along anterior-posterior axis. Points are coloured according to response classification. E. Average baseline subtracted PSTHs of dLGN cells classified as light-responsive recorded either during midday or midnight in response to a 2s light step from darkness at each irradiance (log photons/cm^2^/s, indicated above the panels). E. Peak firing rates during the first 500 ms following light onset increased with irradiance, with irradiance-response relationships differing between circadian phases (extra sum-of-squares F-test: F(3,469) = 4.08, P = 0.007). As in wild-type mice, neurons recorded during the night reached saturation at lower irradiances (see Figure 5F). F. Visual responses were similar at irradiances below (14 log photons/cm²/s) and above (15 log photons/cm²/s) the reported threshold for ReaChR activation, with no effect of opto-responsiveness, light intensity, or their interaction (linear mixed-effects model; opto-responsiveness: t(107.6) = 1.65, P = 0.102; light intensity: t(93) = 0.43, P = 0.670; interaction: t(93) = −0.06, P = 0.950).

## Methods

### Animals, housing and husbandry

All experiments were performed in accordance with the Animals (Scientific Procedures) Act of 1986, and were approved by the University of Manchester Animal Welfare Ethical Review Body and the UK Home Office. 24-hour recordings were performed using male C57BL/6J mice from Charles River aged between 60-120 days. Staircase stimuli experiments were performed using C57BL/6J mice bred inhouse by the University of Manchester Biological Services Facility (males and females balanced across experimental groups) all aged between 60-150 days. For examination of how time of day impacted light-steps from darkness, experiments were performed on 12 C57BL/6J mice (males and females balanced across experimental groups).

For optogenetic control of retinal ganglion cell terminals, we generated Brn3c-Cre x ReaChR mice to drive expression of the light-activated protein ReaChR (Red-shifted Channelrhodopsin) exclusively in neurons expressing Brn3c-Cre (Brn3c-Cre mice kindly shared by Tudor Badea, National Eye Institute, NIH, USA). This labels a heterogenous but consistent subset of retinal ganglion cells (RGCs) which terminate in many retinorecipient regions including the dLGN, but excludes the intrinsically photosensitive ganglion cells (Parmhans et al., 2020). Mice were homozygous for ReaChR but heterozygous for Brn3c-Cre. Mice were aged between 80-240 days, with the numbers of males and females counterbalanced across experimental groups.

All mice were group housed in standard home cages, unless they were recovering from surgery, in which case they were singly housed to avoid cage mates interfering with the implants. All mice were housed in light-tight cabinets to allow for strict control of the lighting environment and had 12hr light: 12 hr dark, under temperature and humidity-controlled environment. Food and water were available *ad libitum*.

### 24 hr recordings from awake animals

#### Recovery Surgery

Animals were anaesthetized using isoflurane (95/5% Oxygen/CO_2_ mix; flow rate: 0.5 - 1.0L/min; 5% for induction and 1% for maintenance). Mice were secured in earbars and kept on a heatmat to maintain a body temperature of 37°C, with lubrithal eye drops to prevents the eyes drying out. 1% EMLA cream was applied topically to the shaven scalp and a subcutaneous injection of buprenorphine (0.05mg/kg) provided analgesia. Following confirmation of a surgical plane of anaesthesia (lack of pedal withdrawal reflex, blink reflex and reduced breathing rate), we used a scalpel to expose the skull surface and drilled a small hole at coordinates above the dLGN (-2.3mm lateral from both Bregma and right from the midline). Larger holes were drilled in the same coordinates on the left hemisphere and at a central point behind lambda to act as anchor points for 2 small screws (M1.6×2.0mm, Precision Tools, UK). Electrodes were coated with DiI (Thermo Fisher Scientific) and lowered into the brain to the depth of the dLGN (∼2.5mm from the surface of the brain), and a grounding wire connected to the screw in the left hemisphere secured with superglue. Light responses were confirmed before UV-curing cement sealed the implant in place. Following recovery, mouse welfare and behaviour were closely monitored for at least a week with further analgesia being administered if necessary.

#### Arena and habituation steps

Mice were acclimatised to both the experimenter and the behavioural arena though short daily handling and habitation sessions (<30mins) for a week prior to the testing day to minimise any stress, with habituation times performed at different times of day to minimise any potential Zeitgeber effects (Tahara et al., 2015). Mice were housed under controlled light / dark schedules for at least 2 weeks prior to test day, with half of the mice on a 12hr reversed light dark schedule (see Fig 1C).

The behavioural arena was adapted from the setup described in Storchi et al (Storchi et al., 2019). In brief, the open field arena (30 x 30cm) was designed to mimic a home cage environment, with bedding, woodchip and wet food provided. The arena was illuminated with infrared lights and a camera (Chameleon 3 from Point Grey) covered with infrared cut-on filters (Edmund Optics) was mounted above the arena to record behaviour in the dark (see Fig 1A).

#### Behavioural protocol

Mice were kept in the dark for at least 18 hours before data collection to control for any impact of light history, but within a time-frame in which circadian time can be well predicted. Start times were staggered such that half the mice began at CT6 (Circadian Time, hours since lights on) and half began at CT18 (see Fig1C). Mice were briefly anaesthetized (<5mins) with 3% isoflurane to attach the head stage under dim red light, placed into the arena and allowed to recover for ∼20 mins. Mice had free exploration of the arena from this point on due to the wireless nature of our recording system and remained in the arena in constant darkness for up to 26 hours. At the end of the experiment, animals were transcardially perfused with 4% PFA and the brains collected for histological verification of electrode placement.

#### Recording spontaneous neural activity and locomotor activity in awake animals

We collected 10 minutes of spontaneous neural activity every 2 hours across the duration of the 24 hours. 16 channel electrodes (Neuronexus; model: A4x4-3mm-50-125-17) were attached to the TBSI W16 wireless acquisition system (Triangular BioSystems; sampling rate = 30 kHz) to allow the recording of multiunit spikes from the dLGN. Corresponding behavioural data was concurrently collected using FlyCapture2 software at a rate of 15 FPS. During epochs between neural data collection, we continued to record locomotor activity but at lower resolution (1FPS).

#### Analysis of Locomotor activity

Videos were analysed offline using the open-source software ezTrack (Pennington et al., 2019), using the Location Tracking Module to track the animal’s location within the arena. The distance travelled is calculated based on the Euclidean distance of the animal’s centre of mass between each subsequent frame. Videos overlaying the estimated centre of mass were generated and manually inspected to ensure correct tracking of the animals. For the high-resolution videos, we applied a 15 frame moving average to smooth the traces and reduce jitter. To determine arousal state, the locomotor data were binned into 10 second intervals and we summed the smoothed distance travelled for each bin. Bins with <5 cm movement per bin were classified as ‘inactive’ and >5 cm as ‘active’ to allow us to assess their general circadian activity patterns.

#### Processing electrophysiology from long term recordings

Continuous data was filtered (4-pole Butterworth, cut-off 250Hz) and spikes were extracted using waveform detection in Offline Sorter V4 (Plexon, US). We removed noise artefacts. For each 10 min epoch, we generated peristimulus time histograms (PSTHs) for each multiunit channel classified as light-responsive during electrode implantation (0.5s pulses of light from darkness). We calculated the time-averaged firing rate for each recording epoch, and in cases where CT6 or CT18 was recorded at both the beginning and the end of the experiment, we calculated the mean of both recordings.

#### Population mean cosinor model

Firing rates were z-scored within each light-responsive multiunit channel across the 24 hr cycle, and then the average z-scored firing rate was calculated at each sampling window calculating the mean across all light-responsive channels. This was used to generate a firing rate activity profile for each mouse across the 24 hours. Missing values (due to equipment malfunction) were imputed using nearest-neighbour linear interpolation. Population-level rhythmicity was assessed using a 24-h population mean cosinor model (R package cosinor2), from which fitted curves, rhythmicity p-values, and percent rhythm (proportion of variance explained by the fitted cosine model) were obtained. For locomotor activity, z-scores were computed on the time-averaged distance travelled per 10 min epoch for each mouse, using the same interpolation and cosinor procedures.

### Acute electrophysiological recordings in anaesthetized mice

#### Surgery and electrode implantation

Surgery was carefully timed so that all data was collected either during specific 4-hour windows ZT5.5-9.5, “midday”; or ZT17.5-21.5, “midnight”. Here, mice were not long-term dark-adapted prior to surgery, and so we refer to experiments in this condition using Zeitgeber Time (ZT; hours since lights on), not Circadian Time (hours since the start of the subjective day) as in the awake electrophysiological experiments. Mice were anaesthetized with 1.4-1.5 g/kg 20 % urethane via intraperitoneal injection, and then a subcutaneous injection of atropine (0.3mg/kg) and sterile saline was delivered to ensure adequate hydration and support breathing. Body temperature was maintained at 37°C using a heatmat. Mice were secured in a stereotaxic frame following confirmation of surgical plane if anaesthesia. Eyes were coated first with atropine (1% in saline, Sigma Aldrich) to dilate the pupils, and then mineral oil (Sigma Aldrich) to protect the cornea for the duration of the recording. We exposed the skull and performed a craniotomy ∼2.3 mm posterior and ∼2.2 mm lateral to Bregma. Recording electrodes coated in DiI were lowered slowly to the target site of the dLGN (∼2.7mm), with the final depth optimised according to location of light responses on the channels. Most experiments were performed using a A4x16-Poly2-5mm-23s-200-177-A64 electrode (Neuronexus), apart from the optogenetic experiments in which case we used a 1X32 OALPpoly3 optrode with connected optical fibre (Neuronexus). The electrode was left in place for 30 min - 1 hr before data collection to allow neural activity to stabilize and to dark adapt the animals. Following completion of recordings, mice were decapitated and the brains were extracted. Brains were incubated for 2-3 days in 4 % PFA at 4 ᵒC before being transferred to 30 % sucrose for cryoprotection.

#### Acute data acquisition

Neural signals were acquired using a Recorder64 system (Plexon), amplified (×3000; or ×3500 for optogenetic experiments), high-pass filtered (300 Hz), and digitized at 40 kHz. Single unit activity was extracted from the multiunit data offline using Kilosort (Pachitariu et al., 2016). Identified single units were exported as ‘virtual tetrodes’ (spike waveforms detected across 4 adjacent channels) and were manually checked in Offline Sorter (Plexon, US). Successfully isolated units displayed a distinct refractory period (> 1.5 ms) in the interspike interval histogram, whereas multiunit activity or noise artefacts were removed.

#### Visual stimuli

For optogenetic experiments, full field visual stimuli were delivered to the eye contralateral to the recording electrode through fibre optic cables generated via a custom light source as described in (Feord et al., 2023). For other experiments, visual stimuli were delivered via liquid light guides and generated via a 4LED light source as described in (Orlowska-Feuer et al., 2023). Neutral density (ND) filter wheels (Thorlabs) allowed for spectrally neutral reductions in 1-log unit steps in light intensity from the brightest light steps (∼15 log photons/s/cm^2^). Short light pulses from darkness were used to identify light-responsive units (2s light steps at the brightest light step, 8s interval of darkness, 10 - 15 repeats). Irradiance response relationships were generated using these same 2s light steps delivered over 6 log units. Fullfield “staircase” stimuli were delivered via the coolLED and consisted of increasing and decreasing steps of background irradiance (9.92, 10.92, 11.92, 12.92, 13.92 and 14.92 log photons/s/cm^2^) with epochs of steady light (30s), temporal white noise (16s) and both contrast chirp and temporal frequency chirps (2s baseline + 8s chirp each) superimposed at each step, as described in (Orlowska-Feuer et al., 2023), see Fig 2C,D. Each full ascension and descension of the staircase was presented 15 times. The contrast chirp stimuli was a sinusoidal modulation which increased from 3% to 98% Michelson contrast centred over the half maximum intensity for that light step. The temporal frequency chirp was an accelerating sinusoidal modulation, maintained at 97% Michelson contrast.

Spatial stimuli were rendered on an LCD display (width: 26.8cm height: 47.4cm; Hanns-G HE225DPB; Taipei, Taiwan) angled at ∼45° from vertical and placed at ∼21cm from the contralateral eye to occupy ∼96° x ∼63° visual angle. Spatial stimuli comprised of a sparse binary noise stimulus (5Hz, square size = 4.2°)(Baden et al., 2016); inverting grating stimuli (1Hz) at spatial frequencies of 0.03 to 1.2 cpd, presented at two phases and four orientations (0°, 45°, 90° and 135°); and a 30s excerpt from naturalistic movie database(Qiu et al., 2021) which was rendered with spectral changes that were designed to recreate UV vs. green chromatic contrast. Stimulus spectra were designed to approximate the activation of each photoreceptor by natural daylight for each species (14.8 MWS-cone opsin effective photons/cm^2^/s; 12.8 SWS-cone effective photons/cm^2^/s; 14.8 rod effective photons/cm^2^/s and 14.7 melanopsin effective photons/cm^2^/s). Intensities were equivalent to those experienced on a cloudy day.

#### Optogenetic Stimuli

Optogenetic stimuli (blue 465 nm 634mW/mm^2^ intensity at the fibre tip via PlexBright LED module, Plexon, USA) were delivered through a 200μm fibre which terminated 100μm above the dorsal-most recording site of the optrode. Light pulses (1 - 500ms delivered every 2s) were controlled through custom written LabView programs (National Instruments, TX, USA), with each flash duration presented 50 times.

### Histology and immunohistochemistry

Brains were sectioned on a freezing sledge microtome (8000 Sledge; Bright Instruments, UK) to collect 40 μm thick coronal slices. Slices were mounted onto slides using Vectashield hardset (H1500, Vector Labs, Peterborough UK) containing DAPI to allow for easier visualisation of tissue cytoarchitecture. The slides were processed through a slide scanner (Laser 2000, UK) in the University of Manchester Bioimaging facility, and the images were acquired using CaseViewer V2.4 software (3DHISTECH, Hungary). The anatomical coordinates of the electrode were estimated based on visualisation of DiI fluorescence, in combination with best-matching coronal panels from the Paxinos and Franklin mouse reference atlas (Paxinos and Franklin, 2003). Projected anatomical locations of each recorded cell was then determined based on the known geometry of the electrode.

### Analysis of acute electrophysiology data

#### Classifying light responses

In all cases, we used a 2s pulse of light (∼15 log photons/s/cm2) from darkness to identify individual light-responsive units. Mean firing rates were calculated during a 2 s pre-stimulus baseline period, a transient ON-response window (0-1 s following light onset), a sustained ON-response window (1-2 s following light onset), and an OFF-response window (0-1 s following light offset). Responses were compared across stimulus repetitions using paired t-tests. Transient and sustained ON responses were assessed relative to baseline firing rate, whereas OFF responses were assessed relative to the sustained response period immediately preceding light offset. Units exhibiting a significant transient increase in firing rate were classified as transient ON, while those exhibiting both transient and sustained increases were classified as sustained ON. Units showing significant transient ON and OFF responses were classified as ON/OFF, and units exhibiting significant decreases in firing rate during either ON-response window were classified as OFF. All remaining units were classified as unclassified.

We constructed PSTHs across repeated presentations of visual stimuli (bin size 50 ms) for individual units, and calculated the mean PSTH across units for average population PSTHs. We used a 5 bin moving average to smooth the data for visualization, but all analyses were conducted on the unsmoothed data. Baseline subtracted PSTHs were calculated by subtracting the mean spike count in the 2s window immediately preceding light presentation and subtracting this from each value in the PSTH.

#### Light steps from darkness

To determine how time of day altered the irradiance response relationships of 2s pulses of light from dark, we calculated the peak change in the average firing rate response in the first 500ms following light onset for each unit. We then averaged across units recorded at each timepoint to obtain population irradiance responses. We fit a 4-parameter logistic (Hill) function to describe the log irradiance response relationships (bottom fixed at 0), comparing a reduced model with parameters shared across timepoints to a full model with parameters specific for each time of day (Top, logEC₅₀, HillSlope).

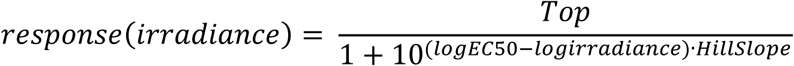

Models were compared via ANOVA (likelihood-ratio test) to assess whether time of day specific parameters improved fit.

#### Analysis of Steady Light steps

To compare how units responded to the ascending vs descending portions of the staircase stimuli, we calculated the mean firing rate for each unit across the 10 - 30s steady light epoch at each irradiance step. For each unit, firing rates were min-max normalized between 0-1 to allow comparison of irradiance response relationships of the population of light-responsive cells when going either up or down the staircase. We fit a Hill function

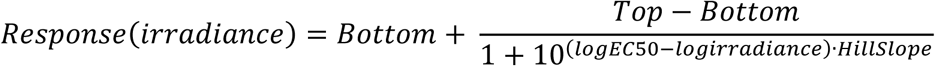

separately for ‘Up’ and ‘Down’ trials (full model) and compared it to a model with shared parameters across directions (reduced model). Nested model comparison (F-test) assessed whether allowing direction-specific parameters provided a significantly better fit.

Data were log-transformed (with a small constant added to avoid zeros) to meet assumptions of normality and homoscedasticity. For comparisons between timepoints, we performed a two-way mixed ANOVA (between-subjects: CT; within-subjects: intensity). Greenhouse-Geisser corrections were applied when sphericity was violated.

#### Contrast sensitivity

Contrast sensitivity was analysed based on the approach described in (Rodgers et al., 2023). For each unit, firing rates were normalised by subtracting the average prestimulus firing rate (0.5s before onset of chirp stim at each light step) and then response amplitudes (maximum firing rate - minimum firing rate) were determined for each 0.5 s bin to generate 16 discrete Michelson contrast values. We fit Naka-Rushton functions to describe the contrast-response relationship, using the four parameter equation

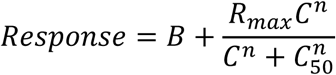

Where B is the baseline offset, *R_max_* is the maximal response amplitude, *n* the slope exponent, and *C_50_* the contrast that produces half the maximum response. For each unit and each irradiance step, the model was fit using nonlinear least-squares regression (R nls, algorithm = “port”) with parameter bounds defined individually from the observed data at that step. Initial parameter values were set from individual minimum/maximum response amplitudes and the mean contrast. We determined the coefficient of determination (R²) and only units that had an R² value of 0.5 or higher were included in subsequent analyses of best fit parameters. We also derived the C_20_ (contrast required to elicit 20% of the modelled maximal response) using these best fit parameters.

We also determined the contrast sensitivity function of the population of light-responsive units recorded at each time of day. To calculate these, each unit’s response amplitude was normalized to its own maximum response across all contrasts and irradiance steps (zero-max normalization). We calculated the average normalized amplitudes at each irradiance step across all units recorded at each time of day, and then the mean population contrast-response curve was fit with the same bounded four-parameter Naka-Rushton model described for single units. The resulting population-level *C_20_*, slope and *R_max_* parameters were used to quantify differences in contrast sensitivity across circadian phase.

#### Temporal frequency tuning

The 8s of sinusoidal modulations in temporal frequency were divided into 25ms bins, which were assigned to one of eight discrete frequency epochs (0-8Hz). For each light-responsive unit and irradiance step, response amplitude was computed for each frequency epoch and normalised to the unit’s maximum response.

Temporal frequency tuning curves were fit with a half-Gaussian model (Grubb and Thompson, 2004; Rodgers et al., 2023) of the form:

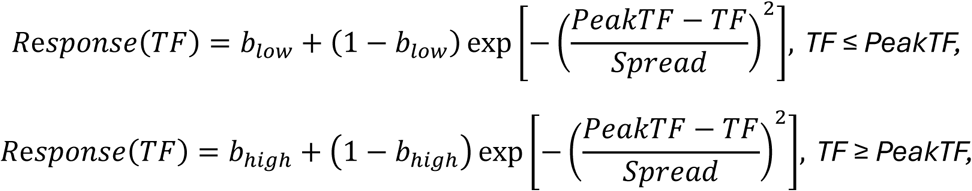

where *PeakTF* is the preferred temporal frequency, *Spread* the Gaussian spread, and *b_low_* and *b_high_* the baseline response levels on the low- and high-frequency sides of the peak. The peak location was systematically fixed to each of the eight candidate temporal frequencies, and the remaining parameters were fit using bounded optimisation. The fit with the lowest residual sum of squares was selected as the final model. Fits with pseudo-R² < 0.5 and with a gaussian spread exactly equal to 1 at brightest lightstep were excluded from subsequent analyses.

We also determined the temporal frequency tuning curves of the population of light-responsive units recorded at each time of day. Each unit’s response amplitudes were normalized using zero-max normalisation, averaging across units within each circadian group and irradiance step, and fitting the same half-Gaussian model to the population mean response.

#### Information Rate

The 16s epoch of temporal white noise was generated by modulating irradiance at 20 Hz with a pseudorandom sequence spanning from dark to twice the irradiance of the steady-light step for each step on the staircase. We pooled together data from the two brightest lightsteps (14 and 15 log photons/s/cm2, “bright”) and the two medium lightsteps (12 and 13 log photons/s/cm2, “mid”) to allow quantification of information transfer rate.

At each level of illumination, we first selected all the cells whose activity was stationary within each level of illumination (trial-to-trial Pearson’s correlation significance test). We then estimated information transfer rate on these cells by applying the same techniques described in (Milosavljevic et al., 2018). Briefly, we first estimated Mutual Information as:

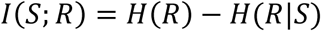

To correct for statistical bias, we applied the information decomposition described in (Panzeri et al., 2007) and applied quadratic extrapolation separately to each term in the decomposition as described in (Strong et al., 1998). We then estimated the information rate (in bits/s) as the slope of I(S; R) as function of time window T (duration = 50, 100, 150, and 200 ms) given a fixed temporal sub-window dt = 50ms. Finally, coding efficiency was estimated by dividing information rate by the average firing rate of each cell.

Log-transformed firing rate and information rates were analysed using linear mixed-effects models (lme4, R), with time-of-day and irradiance as fixed effects and individual units as a random intercept. P-values were computed using Kenward-Roger approximation.

An equivalent approach was used to analyse data from the naturalistic movie stimulus, which was only presented at one stimulus intensity, and x30 repeats.

#### Receptive Field mapping

The spatio-temporal receptive field was derived for each unit by generating the spike triggered average (STA) of responses to a sparse binary noise stimulus (5Hz, square size = 4.2°), as described previously (Allen et al., 2025). The separable spatial and temporal components where then extracted from the raw STA matrix by finding the signal peak. RF locations and sizes were then generated by fitting spatial receptive fields with 2D Gaussian function (using lsqcurvefit function, MATLAB). The receptive field size for individual cells was the average of the standard deviation of Gaussians fitted to each dimension.

#### Spatial frequency tuning and spatial summation

To assay changes in spatial frequency tuning, inverting gratings (Michelson contrast between dark and light bars = 98%) were presented in 4 orientations at two phases (phase shifted 90°), at 5 different spatial frequencies (0.03 to 1.2 cpd) at 2Hz. For each unit, response amplitudes were quantified (relative to pre-stimulus firing) for each phase/orientation combination for each spatial frequency to determine the optimal stimulus (R_pref_). R_orth_ was the response to stimuli presented at 90° to the preferred orientation. Continuous firing rates during the stimulus presentation were then Fourier analysed to extract amplitudes of the first and second harmonic components (F1 and F2, at 1Hz and 2Hz), at the preferred and null (90° phase shifted) stimulus.

These data were also used to quantify an orientation selectivity index in responding units. OSI was calculated as the ratio of (R_pref_-R_orth_)/(R_pref_+R_orth_) at the preferred spatial frequency.

#### Classifying opto-responsive units

For experiments using optogenetic stimuli, PSTHs were generated with bin sizes of 5ms. For each unit, we quantified whether the change in spike rate in the 50 ms immediately following optogenetic stimulation (relative to that in the preceding 100 ms) was significantly greater than zero (one sample t-tests; P < 0.05). Cells were classified as opto-responsive if they showed a statistically significant excitatory response to at least 2 stimulus durations. Responses for individual cells were baseline corrected by subtracting the mean spike count in the 100 ms window preceding stimulation.

#### Opto-pulse response curves

For quantification of response amplitude across stimulus durations, we defined ‘Peak’ response as maximum the average change in spike rate occurring in the initial 50ms window following opto-pulse onset. Pulse duration-peak response curves were fit to describe the relationship using non-linear regression with a four parameter logistic function, with bottom asymptote constrained to 0. Stimulus duration was log-transformed before fitting. This function took the form :

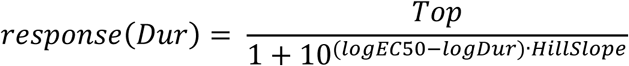

We compared two models; the first allowed separate parameters (Top, logEC50 and Hillslope) for each time of day, whereas the second fit a single set of parameters across both timepoints. Both models were fit using biologically informed starting values derived from the empirical maxima and average log stimulus durations. Improvement in model fit was assessed using likelihood ratio tests via ANOVA.

#### Data presentation

All graphs are presented with mean ± SEM unless otherwise stated. For all data visualizations, experiments performed during midday are in orange, midnight in blue.

